# Reprogramming early cortical astroglia into neurons with hallmarks of fast-spiking parvalbumin-positive interneurons by phospho-site deficient Ascl1

**DOI:** 10.1101/2023.11.03.565289

**Authors:** Nicolás Marichal, Sophie Péron, Ana Beltran Arranz, Chiara Galante, Franciele Franco Scarante, Rebecca Wiffen, Carol Schuurmans, Marisa Karow, Sergio Gascón, Benedikt Berninger

**Author notes:** These authors contributed equally to this work (Nicolás Marichal, Sophie Péron, Ana Beltran Arranz).

## Abstract

Cellular reprogramming of mammalian glia to an induced neuronal fate holds potential for restoring diseased brain circuits. While the proneural factor Ascl1 is widely used for neuronal reprogramming, in the early postnatal mouse cortex Ascl1 fails to induce glia-to-neuron conversion, instead promoting proliferation of oligodendrocyte progenitor cells (OPC). Since Ascl1 activity is post-translationally regulated, here we investigated the consequences of mutating six serine phospho-acceptor sites to alanine (Ascl1SA6) on lineage reprogramming *in vivo*. Ascl1SA6 exhibited increased neurogenic activity in glia of the early postnatal mouse cortex, an effect enhanced by co-expression of Bcl2. Genetic fate-mapping revealed that most induced neurons originated from astrocytes while only a few derived from OPCs. Intriguingly, many Ascl1SA6/Bcl2-induced neurons expressed parvalbumin and were capable of high-frequency action potential firing. Our study demonstrates authentic conversion of astroglia into neurons featuring subclass hallmarks of cortical interneurons, advancing our scope of engineering neuronal fates in the brain.

## Introduction

Changing the identity of a terminally differentiated cell into another desired cell type via lineage reprogramming is an emerging experimental strategy for tissue repair in organs with little or no endogenous stem cell activity (*1*). In the context of the CNS, the generation of induced neurons (iNs) from other resident cell types, such as glial cells, opens new avenues for the remodeling and restoration of diseased brain circuits (*2*). Following pioneering studies demonstrating the possibility of generating iNs from primary glia *in vitro* (*3*, *4*), proof-of-principle studies have since provided evidence for glia-to-neuron conversion in various brain areas *in vivo* through ectopic expression of proneural transcription factors (*5–9*). More recently, however, important concerns about the authenticity and efficiency of *in vivo* glia-to-neuron conversion have been raised since many studies lacked effective lineage tracing to demonstrate the glial origin of iNs (*10–12*). Indeed, subsequent efforts that scrutinized the origin of putative iNs uncovered inadvertent labeling of endogenous neurons that could not be traced back to a glial origin (*10*). This emphasizes the importance of robust genetic fate mapping to unambiguously determine the glial origin of iNs (*13–15*).

The basic helix-loop-helix (bHLH) transcription factor achaete-scute complex-like 1 (Ascl1) is a key regulator of neural fate decisions ranging from cellular proliferation and cell cycle exit to neural fate specification in the embryonic and adult nervous system (*16–20*). Given its neurogenic activity, Ascl1 has been widely and successfully used for converting somatic cells such as fibroblasts, glia and other cell types into iNs *in vitro,* either alone or in combination with other transcription factors (*4*, *21*, *22*). However, the neurogenic activity of Ascl1 is highly cell- and context-dependent. For instance, on its own, Asc1 failed to convert glia in the early postnatal cortex into neurons *in vivo* and instead promoted the proliferation of oligodendrocyte progenitor cells (OPCs) (*23*). Likewise, when expressed without other factors, Ascl1 reprogrammed reactive glia in the lesioned adult cortex or the epileptic hippocampus into iNs only inefficiently (*5*, *7*). While several studies have highlighted the capacity of Ascl1 to act as a pioneer transcription factor (*24*, *25*), the high degree of variability of Ascl1 effects *in vivo* is suggestive of additional layers of regulation that may hinder its neurogenic activity. Indeed, Ascl1 activity has been shown to be regulated by post-translational modifications, including phosphorylation (*26–29*). Interestingly, preventing phosphorylation-dependent regulation of Ascl1 activity by mutating serine residues of six serine-proline motifs to alanine (Ascl1SA6), has been found to increase its neurogenic activity in the embryonic cerebral cortex (*26*). This finding has led us to hypothesize that employing the Ascl1SA6 mutant variant as a reprogramming factor could enhance glia-to-neuron conversion *in vivo*.

Here we demonstrate that retroviral-mediated overexpression of Ascl1SA6 in glia undergoing proliferative expansion in the early postnatal mouse cerebral cortex has enhanced neurogenic capacity compared to that of wildtype (wt) Ascl1 *in vivo.* Furthermore, we found that co-expression of Ascl1SA6 with B-cell lymphoma 2 (Bcl2), previously demonstrated to enhance iN survival (*6*), can more efficiently reprogram postnatal cortical glia into iNs. Through genetic fate mapping, we show that iNs originate predominantly from astrocytes and to a much lesser degree from OPCs. Intriguingly, we also found that Ascl1SA6/Bcl2-induced neurons frequently exhibit hallmarks of fast-spiking (FS) parvalbumin (PV)-positive interneurons, an important subclass of GABAergic interneurons in the mammalian cerebral cortex.

## Results

### A phospho-site mutant of Ascl1 exhibits enhanced neuronal reprogramming activity *in vivo*

We have recently established an experimental model to explore glia-to-neuron conversion in the early postnatal mouse cerebral cortex (*23*, *30*), in which MMLV (Moloney murine leukemia virus) retroviruses enable efficient targeting of both astrocytes and oligodendrocyte progenitor cells (OPCs) undergoing proliferative expansion (*31*, *32*). Given that we had previously observed that Ascl1 failed to induce substantial neurogenesis (*23*), here we wondered whether utilizing a more neurogenic phospho-site mutant Ascl1 (Ascl1SA6) (*26*, *27*), in which six serine-proline sites had been altered to alanine-proline exhibited improved neuronal reprogramming (Fig. S1A). To address this question, we compared the effects of Ascl1- or Ascl1SA6-encoding retroviruses upon injection in the mouse cerebral cortex at postnatal day 5 (P5). When analyzed at 12 days post injection (dpi), a time point when previous studies had observed successful glia-to-neuron conversion by other neurogenic transcription factors (*5*, *6*, *30*), we found that Ascl1 elicited very limited expression of immature and mature neuronal markers doublecortin (Dcx) and NeuN, respectively, in transduced cells, which mostly retained glial morphology (Fig. S1B, C). In sharp contrast, MMLV retrovirus-mediated expression of Ascl1SA6 induced neuronal marker expression (Dcx, NeuN, or both) in almost half of the transduced cells (Fig S1B, C). These data suggest that the activity of Ascl1 in proliferative glia is subject to post-translational regulation and that the phospho-site mutant is more powerful in inducing neuronal fate conversion from glia than wt Ascl1.

Glial cells undergoing reprogramming by forced expression of proneural transcription factors are susceptible to cell death via ferroptosis, which can be mitigated by co-expression of B-cell lymphoma 2 (Bcl2) (*6*). Thus, we next examined the effect of Bcl2 co-expression alongside Ascl1 or Ascl1SA6 (Fig. 1A). Cells transduced with Bcl2 alone (GFP+) retained glial identity (Fig. 1B). While Ascl1 expression alone showed negligible lineage conversion (Fig S1A), when co-expressed with Bcl2 we observed a significant neurogenic response, as determined by Dcx and NeuN immunoreactivity in double-transduced cells (GFP+/RFP+) at 12 dpi (Fig. 1B, D; Fig. S2). Neuronal fate conversion was further enhanced when Bcl2 was co-expressed with Ascl1SA6, resulting in higher levels of NeuN expression and the appearance of cells with clear neuronal morphologies (Fig. 1B, D; Fig. S2). Twenty-eight days after retroviral injection (28 dpi), the vast majority of Ascl1/Bcl2 iNs did not express NeuN and maintained glial morphology, while around 80% of Ascl1SA6/Bcl2 iNs expressed NeuN (Fig. 1C, D). Thus, the enhanced neurogenic activity of the phospho-site mutant Ascl1SA6 versus wt Ascl1 is further potentiated when combined with Bcl2.

**Fig. 1.**
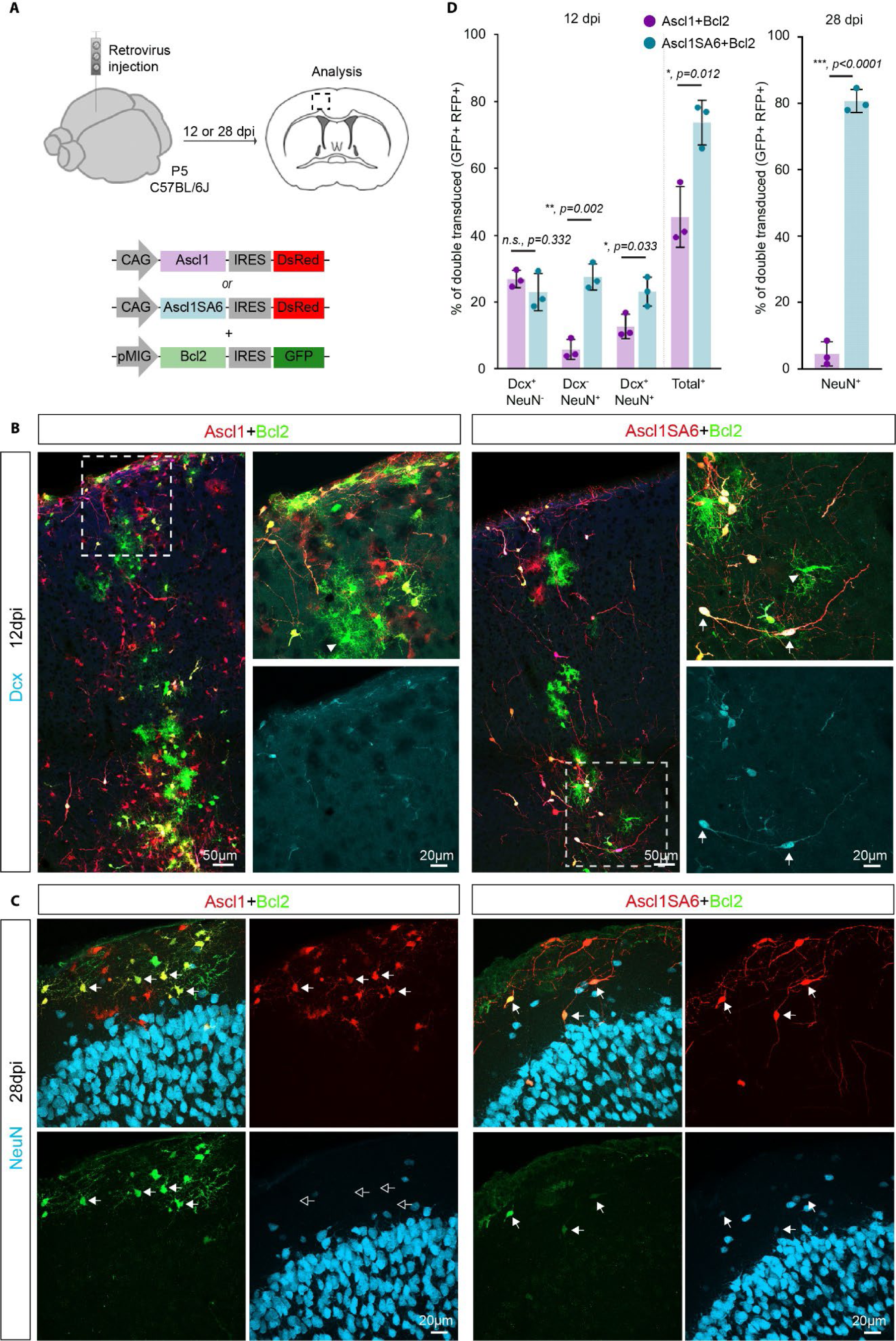
Co-expression of Ascl1SA6 and Bcl2 efficiently reprograms postnatal cortical glia into iNs. (A) Experimental design. Retroviral constructs encoding for Ascl1 or Ascl1SA6 and Bcl2 were injected in the cortex of wt (C57BL/6J) mice at P5. Immunohistochemical analysis was performed at 12 or 28 dpi. (B) Low-magnification image showing Ascl1 or Ascl1SA6/Bcl2-transduced cells at 12 dpi. High-magnification images from insets (white boxes) depicting Dcx expression in co-transduced cells. Notice the acquisition of conspicuous neuronal morphology in Ascl1SA6/Bcl2 iNs (arrows). Cells transduced with Bcl2 only (GFP+ only) retained glial morphology (arrowheads). (C) At 28 dpi, the vast majority of Ascl1SA6/Bcl2 iNs expressed NeuN, while Ascl1/Bcl2 iNs did not express this neuronal marker and maintained glial morphology. Empty arrows indicate marker-negative cells. (D) Proportion of double-transduced cells expressing Dcx, NeuN or both neuronal markers at 12 dpi (left graph), and double-transduced cells expressing NeuN (right graph) at 28 dpi. 26.9 ± 2.6% Dcx only, 5.8 ± 3% NeuN only, 12.7 ± 3.6% Dcx and NeuN, 2380 cells, n = 3 mice for Ascl1 /Bcl2; 23 ± 5.5% Dcx only, 27.5 ± 3.9% NeuN only, 23.2 ± 4.3% Dcx and NeuN, 836 cells, n = 3 animals for Ascl1SA6/Bcl2 at 12 dpi. 4.5 ± 3.6% NeuN, 499 cells, n = 3 mice for Ascl1/Bcl2; 80.7 ± 3.5% NeuN, 157 cells, n = 3 mice for Ascl1SA6/Bcl2 at 28 dpi. Data shown as mean ± SD. Two-tailed Student’s unpaired t-test in (D).

To ascertain the specificity of the MMLV retroviral vectors in stably transducing only proliferating glial cells and rule out unintended labeling of endogenous post-mitotic neurons (*10*), we injected thymidine analogue bromodeoxyuridine (BrdU) to label proliferating cells on the day of, and day following, MMLV retrovirus injection (Fig. S3A). As expected, we found that a large proportion of transduced cells were BrdU-positive (Fig. S3B, C). Importantly, we found that the vast majority (∼ 80%) of retroviral transduced cells that had acquired a neuronal morphology were BrdU-positive (Fig. S3B, C) and co-expressed NeuN (Fig. S3D), indicating that iNs derived from cells that had proliferated during this stage of postnatal cortical development. Interestingly, we consistently observed that the level of the BrdU signal was lower in cells retaining glial morphology, possibly due to dilution of the BrdU signal as a result of continued proliferation (Fig. S3E-G). These data provide indirect evidence that lineage conversion from glia to neuron occurs in the absence of significant cell division, as iNs maintain significantly higher BrdU signal levels as compared to non-reprogrammed glia (Fig. S3E-G).

### Astroglial origin of Ascl1SA6/Bcl2 iNs

Several glial cell types undergo local proliferation in the early postnatal cortex and could, therefore, be targeted by our retroviral constructs (*31–33*), raising the question of the cellular origins of iNs following Ascl1SA6 and Bcl2 mediated lineage conversion in early postnatal cortex. To address this question, we combined reprogramming with Ascl1SA6 and Bcl2 together with genetic fate mapping. Astrocytes were fate-mapped using Aldh1l1-CreERT2/RCE:loxP transgenic mice (*34*), in which Cre-dependent recombination of CAG-boosted EGFP expression was induced by administration of tamoxifen in the days preceding viral injection (Fig. 2A). To allow for tracing EGFP-positive cells expressing the two reprogramming factors, we designed a single retroviral vector encoding for both Ascl1SA6 and Bcl2 (Ascl1SA6-Bcl2) and the DsRed reporter (Fig. 2A), which exhibited a similar reprogramming efficacy as the individual vectors when injected in wildtype mice (Fig. S4A-C). Tamoxifen-treated Aldh1l1-CreERT2/RCE:loxP mice were injected with the Ascl1SA6-Bcl2 encoding retrovirus at P5 and the identity of fate-mapped cells was analyzed by immunostaining for GFP (identifying cells of astroglial origin), DsRed (identifying transduced cells) and Dcx or NeuN (identifying iNs) at 12 dpi (Fig. 2A). We found that more than two-thirds of the DsRed-positive transduced cells also expressed EGFP (i.e., retrovirus-targeted cells of astroglial origin, Fig. 2B-E white and yellow sections, large pie charts). Of these transduced astroglial cells, a high proportion was immunoreactive for Dcx (Fig. 2B, D) and NeuN (Fig. 2C, E). Importantly, of all cells classified as iNs (Fig. 2D, E white and pink sections, large pie charts) 80 % could be lineage-traced to astrocytes (Fig. 2D, E white sections, small pie charts). To independently confirm these results, we performed lineage tracing in mGFAP-Cre/ RCE:loxP mice in which astrocytes are stably labelled with EGFP (*35*) (Fig. S5A). Similarly, we found that a substantial proportion of Dcx-positive and NeuN-positive transduced cells were also EGFP-positive (Fig. S5B-E). These results unambiguously confirmed the astroglial origin of a substantial proportion of Ascl1SA6-Bcl2 iNs.

**Fig. 2.**
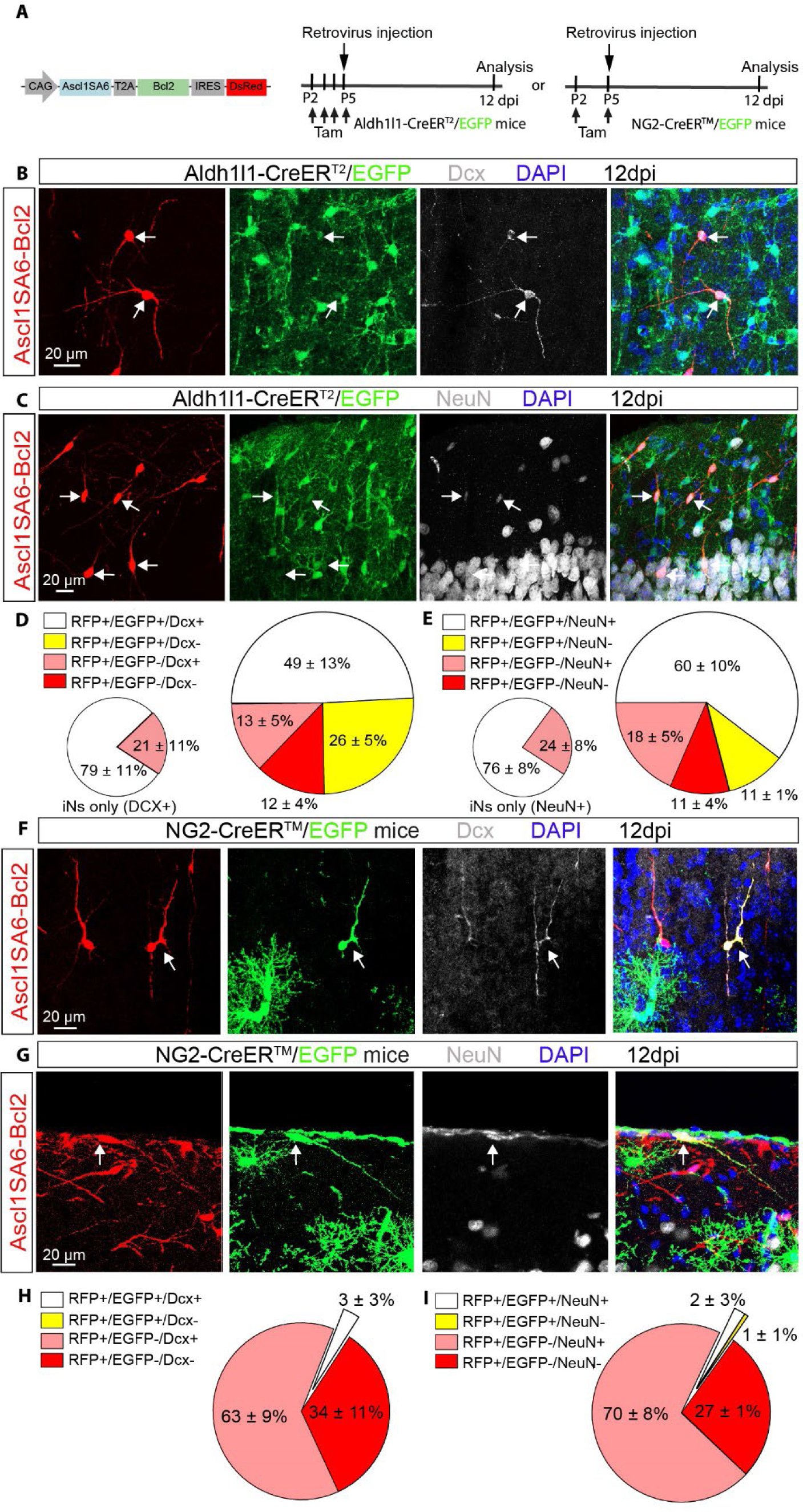
Astroglial origin of the vast majority of Ascl1SA6-Bcl2-derived iNs. (A) Experimental design. Retroviral constructs encoding for Ascl1SA6-Bcl2 were injected in the cortex of Aldh1l1-CreERT2/RCE:loxP or NG2-CreERTM/RCE:loxP transgenic mice at P5. Mice received subcutaneous injection of tamoxifen before retroviral injection to induce Cre-mediated recombination and achieve irreversible labelling of astrocytes or oligodendrocyte precursor cells with EGFP. The proportion of cells with astroglial or oligodendroglial origin was analyzed at 12 dpi. (B-C) Ascl1SA6-Bcl2-derived iNs expressing Dcx (B) or NeuN (C) and GFP (arrows) in the Aldh1l1-CreERT2/RCE:loxP mice, demonstrating their astroglial origin. (D-E) Large pie charts showing the relative number of transduced cells (DsRed+) co-expressing EGFP and/or Dcx (D) and/or NeuN (E), or no other marker than the DsRed in Aldh1l1-CreERT2/RCE:loxP mice at 12 dpi. 523 cells, n = 3 mice for Dcx analysis; 473 cells, n = 3 mice for NeuN analysis. Small pie charts include only those DsRed+ cells that also expressed a neuronal marker (Dcx or NeuN); white sections, lineage traced (EGFP+); pink sections, no lineage tracing (EGFP-). (F-G) Ascl1SA6-Bcl2-derived iNs co-expressing Dcx (B) or NeuN (C) as well as the reporter genes DsRed and GFP in NG2-CreER^TM^/RCE:loxP mice at 12 dpi. (D-E) Pie charts showing the relative number of transduced cells (DsRed+) co-expressing EGFP and/or Dcx (D) and/or NeuN (E), or no other marker than DsRed in NG2-CreER^TM^/RCE:loxP mice at 12 dpi. 416 cells, n = 3 mice for Dcx analysis; 169 cells, n = 3 mice for NeuN analysis. Data shown as mean ± SD.

OPCs also sustain high proliferation in the postnatal cortex (*31*) and are transduced by retroviruses injected in the P5 postnatal cortex (*23*). We therefore tested next whether cells of the oligodendroglial lineage were also converted by Ascl1SA6-Bcl2 into Dcx-positive or NeuN-positive iNs. To this end, genetic fate mapping was performed in NG2-CreERTM/RCE:loxP transgenic mice, in which cells with an active *Cspg4* locus encoding neuron-glial antigen 2 (NG2) are stably labelled by EGFP reporter expression upon tamoxifen injection (*36*) (Fig. 2A). Surprisingly, we found that only approximately 3% of DsRed-positive transduced cells also expressed EGFP and Dcx or NeuN (Fig. 2F-I), suggesting that a very small fraction of the Ascl1SA6-Bcl2 iNs were of oligodendroglial origin. To validate that MMLV retrovirus-transduced OPCs could be effectively lineage traced, we injected a control MMLV retrovirus (Fig. S5F). We found that 22% of all transduced cells could be lineage traced to the oligodendroglial lineage (Fig. S5G-H), consistent with a previous characterization of MMLV retrovirus targeted glial population (*23*). As expected, transduced cells that were not fate-mapped were mostly Sox9-positive astrocytes (Fig. S5G-H). Altogether, our genetic fate mapping experiments demonstrate the authenticity of Ascl1SA6-Bcl2 glia-to-neuron lineage conversion in the early postnatal cortex, with astrocytes being the predominant cellular origins for observed iNs. Despite our retroviral vectors being able to successfully transduce cortical OPCs, these contribute only a very small fraction of iNs.

### Parvalbumin expression in Ascl1SA6/Bcl2 iNs

A central aim of glia-to-neuron conversion is to identify reprogramming factors that allow for the generation of neurons with subtype-specific features *in vivo*. Ascl1 is known to be a pioneer transcription factor involved in GABAergic neuronal fate specification at early embryonic stages (*37–39*). At 28 dpi, we detected GABA immunoreactivity in a very small fraction of Ascl1/Bcl2 iNs. In contrast, approximately 25% of Ascl1SA6/Bcl2 iNs expressed this marker (Fig. 3A, B), consistent with a role of unphosphorylated Ascl1 in GABAergic neurogenesis (*26*). To further characterize these iNs, we next examined whether they had acquired hallmarks of main subclasses of cortical interneurons such as expression of the neuropeptide somatostatin (SST) or the calcium-binding protein parvalbumin (PV) (*40*). At 28 dpi, SST mRNA was found neither in Ascl1/Bcl2 iNs nor in AsclSA6/Bcl2 iNs (Fig. 3C). Likewise, virtually none of the Ascl1/Bcl2 transduced cells expressed PV. Remarkably, however, approximately 20% of Ascl1SA6/Bcl2 iNs acquired PV immunoreactivity at 28 dpi (Fig. 3D, E). Further, smFISH revealed that Ascl1SA6/Bcl2 iNs expressed PV mRNA already at 12 dpi (Fig. 3F, G), albeit at lower levels than endogenous PV cortical interneurons (Fig. 3G). Taken together, these results show that overexpression of Ascl1SA6 and Bcl2 can induce the conversion of postnatal cortical glia into iNs expressing both GABA and PV.

**Fig. 3.**
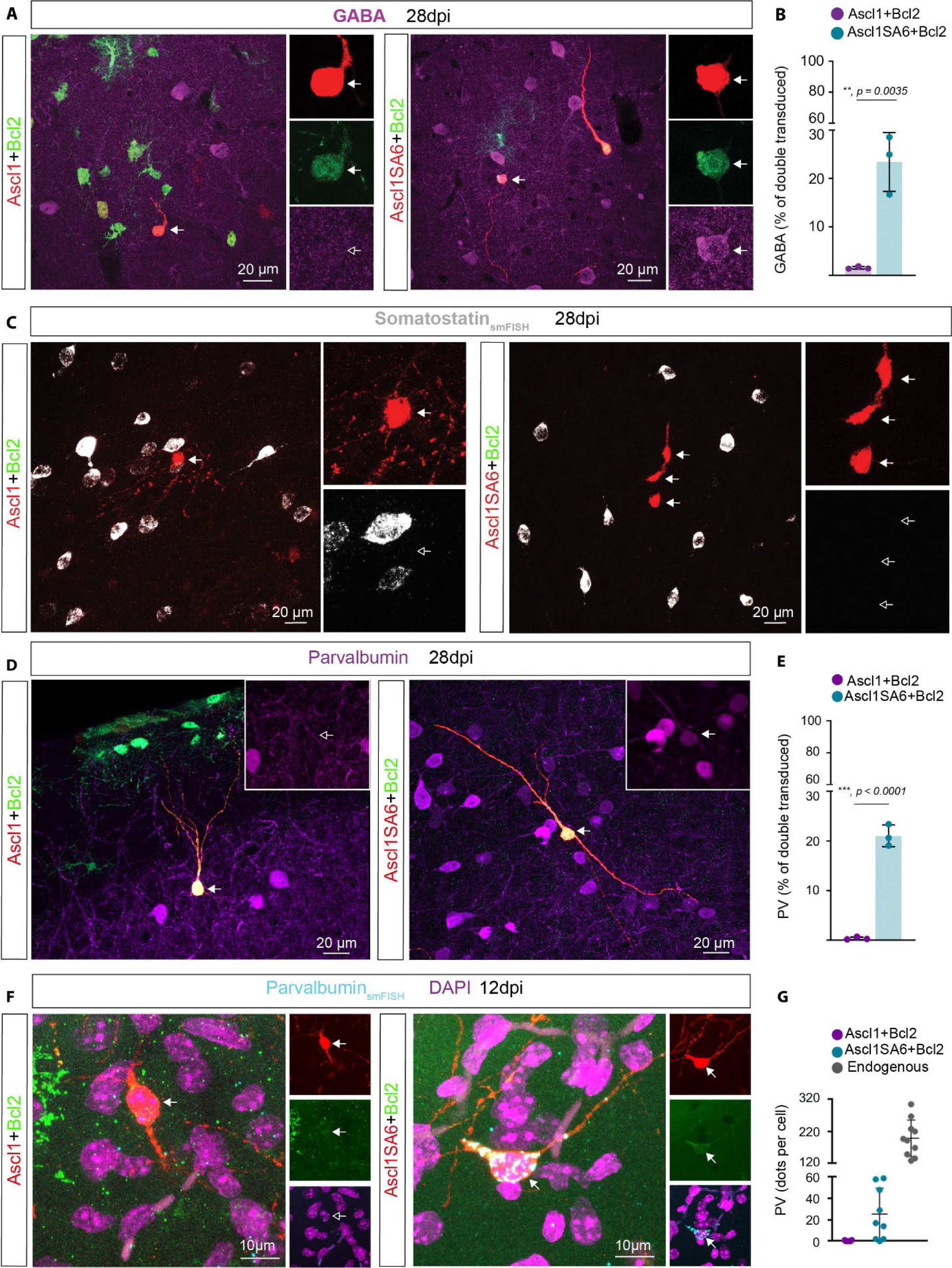
Forced co-expression of Ascl1SA6 and Bcl2 converts postnatal cortical glia into GABA and PV-expressing iNs. (A) Ascl1/Bcl2-transduced cells do not express GABA (left panel), in contrast to the acquisition of GABA expression in Ascl1SA6/Bcl2 iNs (right panel), 28 dpi. Arrows indicate iNs lacking GABA (inset, empty arrow) or containing GABA (inset, filled arrow). (B) Proportion of double-transduced cells expressing GABA following Ascl1/Bcl2 or Ascl1SA6/Bcl2 reprogramming at 28 dpi. 1.6 ± 0.3%, 634 cells, n = 3 mice for Ascl1/Bcl2, 23.4 ± 6.1%, 78 cells, n = 3 mice for Ascl1SA6/Bcl2. Data shown as mean ± SD. (C) Ascl1/Bcl2 and Ascl1SA6/Bcl2 iNs lack SST mRNA. Empty arrows highlight lack of SST expression in iNs (insets). (D) Analysis of PV expression in Ascl1/Bcl2-transduced cells (left panel) and Ascl1SA6/Bcl2 iNs at 28 dpi. Arrows indicate iNs lacking PV (inset, empty arrow) or containing PV (inset, filled arrow). (E) Proportion of double-transduced cells expressing PV at 28 dpi. 0.4 ± 0.3%, 2995 cells, n = 3 mice for Ascl1/Bcl2; 21.0 ± 2.2%, 208 cells, n = 3 mice for Ascl1SA6/Bcl2. Data shown as mean ± SD. (F) Ascl1/Bcl2-transduced cells (left panel) lack PV mRNA, while Ascl1SA6/Bcl2 iNs (right panel) express PV mRNA already at 12 dpi. Empty arrows indicate marker-negative cells. (G) Quantification of PV mRNA transcripts expressed as the total number of dots detected per individual cell in Ascl1/Bcl2, Ascl1SA6/Bcl2-transduced cells and endogenous surrounding PV interneurons. Each dot represents one cell. 7 cells, n = 3 mice for Ascl1/Bcl2; 9 cells, n = 3 mice for Ascl1SA6/Bcl2 cells; 10 cells, n = 3 mice for endogenous neurons. Data shown as mean ± SD. Two-tailed Student’s unpaired t-test in (B) and (E).

### Fast-spiking phenotype of Ascl1SA6/Bcl2 iNs

PV-positive cortical interneurons show characteristic electrophysiological properties including fast-spiking (FS) firing (*40*, *41*). Given the expression of GABA and parvalbumin in our Ascl1SA6/Bcl2 iNs, we next asked whether they acquired functional properties reminiscent of FS/PV interneurons by performing patch-clamp recordings in acute *ex vivo* cortical slices at 28 dpi (Fig. 4A, C). While Ascl1/Bcl2 iNs, mostly retained glial-like membrane properties, Ascl1SA6/Bcl2 iNs showed significantly higher input resistances, and exhibited an overall tendency towards more negative resting membrane potentials and greater membrane capacitances, which were suggestive of an immature neuronal phenotype (Fig. S7A-C). Consistent with this, Ascl1/Bcl2 cells had the ability to generate only a single spikelet followed by non-regenerative membrane potential oscillations in response to sustained depolarization (Fig. 4B). This single spikelet was blocked by tetrodotoxin (Fig. 4B), confirming that it was mediated by voltage-gated Na^+^ channels. Only one Ascl1/Bcl2 cell out of 13 cells recorded showed passive responses to depolarizing current pulses.

**Fig. 4.**
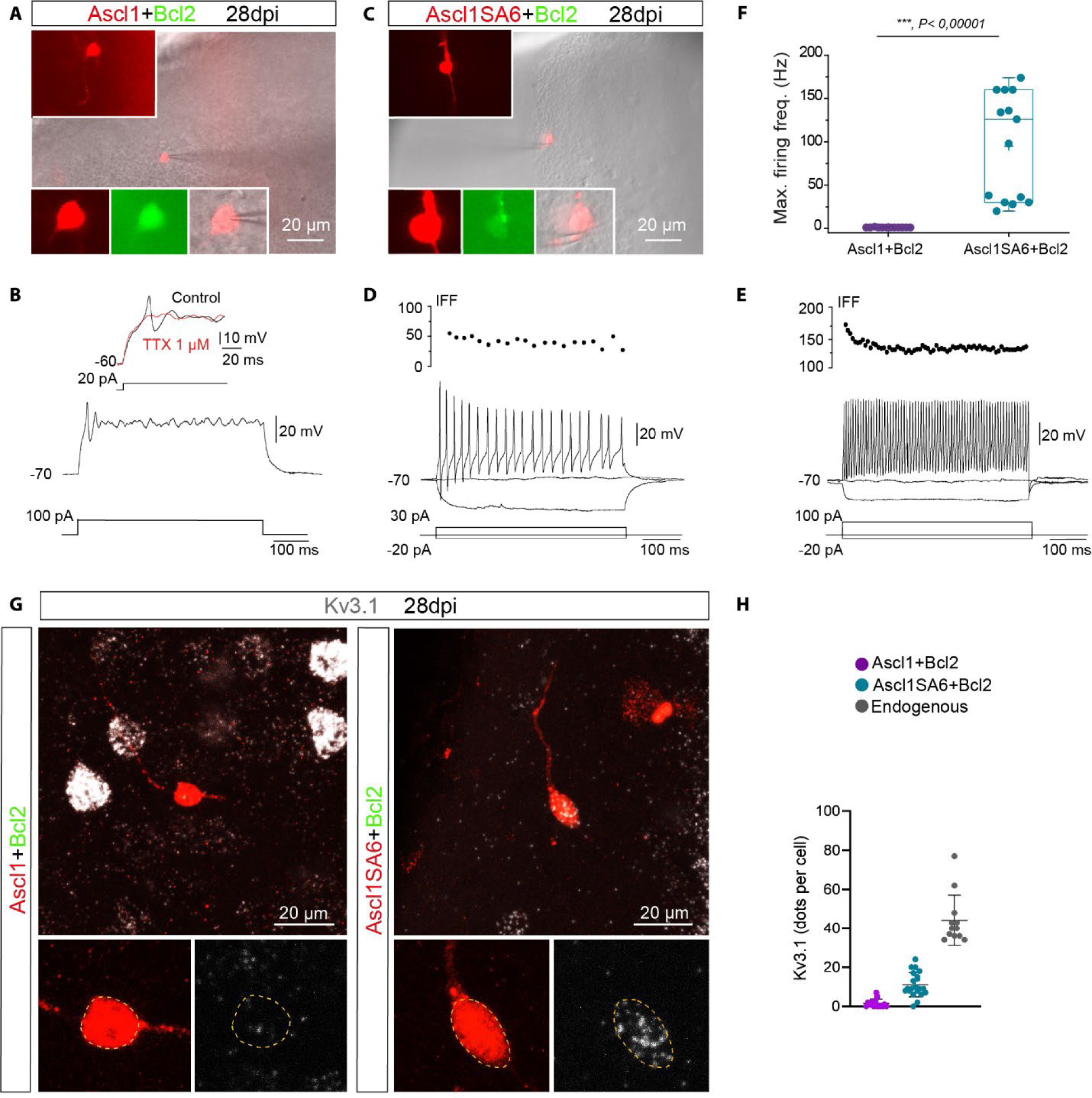
Induced neurons generated by co-expression of Ascl1SA6 and Bcl2 acquired functional properties of fast-spiking interneurons. (A) Example of recorded Ascl1/Bcl2 co-transduced cell visualized in an acute brain slice. Insets show DsRed expression depicting the morphology of the cell (upper inset) and colocalization of DsRed and GFP in the soma (lower insets). (B) Ascl1/Bcl2 recorded cell showing a single small spike in response to depolarizing current injection (12 of 13 recorded cells, n = 5 mice). The single spike was sensitive to TTX (3 of 3 cells), indicating that was mediated by voltage-gated Na+ channels. (C) Recorded Ascl1SA6/Bcl2 co-transduced cell visualized in an acute brain slice. Insets similar as in (A). (D) Ascl1SA6/Bcl2 iN showing repetitive action potential with an instantaneous firing frequency (IFF) in the range of 50 Hz in response to depolarizing current injection (6 of 14 recorded cells, n = 8 mice). (E) Ascl1SA6/Bcl2 iN showing sustained high IFF (>100 Hz, 8 of 14 recorded cells, n = 8 mice). (F) Maximum firing frequency values in recorded Ascl1/Bcl2 and Ascl1SA6/Bcl2 iNs. Each dot represents one cell. Data shown with box and whisker plots, which give the mean (+), median, 25th and 75th percentiles and range. (G) iNs expressing mRNA of the voltage-gated K+ Kv3.1. (H) Quantification of Kv3.1 mRNA transcripts expressed as the total number of dots detected in individual cells in Ascl1/Bcl2, Ascl1SA6/Bcl2-tranduced cells, and endogenous surrounding interneurons. Each dot represents one cell. 15 cells, n = 3 mice for Ascl1/Bcl2-transduced cells; 21 cells, n = 3 mice for Ascl1SA6/Bcl2-transduced cells; 12 cells, n = 3 mice for endogenous neurons. Two-tailed Mann Whitney test in (F).

In sharp contrast, all Ascl1SA6/Bcl2 cells recorded (n=14 cells) were able to repetitively fire well-developed action potentials in response to depolarization (Fig. 4C-E). About half of them generated sustained trains of spikes with an instantaneous firing frequency in the range of 50 Hz (Fig. 4D). Remarkably, the other half of the recorded cells exhibited sustained high-frequency firing, reaching frequencies above 150 Hz with a small drop in frequency firing during a period of constant activation (i.e., adaptation) (Fig 4E, F) - two distinctive features of endogenous cortical FS interneurons (*42*, *43*). Most Ascl1SA6/Bcl2 iNs exhibited a narrow action potential that included a well-developed afterhyperpolarization (AHP) in response to 10-ms somatic current injection (Fig. S6A), resembling action potential waveforms of endogenous FS interneurons (*43*). As expected, the amplitude of the action potential and the AHP of Ascl1SA6/Bcl2 iNs were significantly higher, and the action potential half-width was significantly shorter than their counterparts in Ascl1/Bcl2 iNs (Fig. S6B-D). We also evaluated whether iNs became integrated into the host circuitry by monitoring spontaneous postsynaptic currents. Consistent with the immature functional phenotype of Ascl1/Bcl2 iNs, only a small proportion of these cells exhibited spontaneous synaptic currents (Fig. S7D, F, G). In contrast, the vast majority of Ascl1SA6/Bcl2 iNs exhibited postsynaptic currents (Fig. S7E, F, G) that were blocked by the AMPA/kainate receptor blocker CNQX (Fig S7H). Interestingly, while the number of iNs receiving synaptic input were fewer in the Ascl1/Bcl2 compared to AScl1SA6/Bcl2 group (Ascl1/Bcl2: 4/13 cells vs Ascl1SA6/Bcl2: 11/14 cells), the inputs did not differ significantly in frequency and amplitude between the two cohorts (Fig. S7F, G). Since we observed FS properties in iNs, we analyzed the expression of the delayed rectifying potassium channel Kv3.1 in iNs. Kv3.1 expression is mainly restricted to FS cortical interneurons and is critical for FS properties by providing rapid membrane repolarization after an action potential (*44*). Accordingly, we found that most of the Ascl1SA6/Bcl2 iNs expressed Kv3.1 mRNA and protein (Fig 4G; Fig. S6E), albeit at lower levels than endogenous interneurons (Fig 4H, Fig. S6E). Overall, these data demonstrate that Ascl1SA6/Bcl2 can convert cortical glia into iNs that acquire functional hallmarks of FS interneurons.

### Induced fast-spiking and parvalbumin-expressing iNs in cortical layer I

In the mouse cortex, FS and PV-expressing interneurons locate to layers II-VI but are absent from layer I (*41*) (Fig. 5A). Given that Ascl1SA6/Bcl2 can induce the expression of PV in iNs, we next addressed whether these reprogramming factors could induce PV expression in an ectopic location. Interestingly, we found PV-positive Ascl1SA6/Bcl2 iNs located in cortical layer I (Fig. 5B, C). At 28 dpi, the relative proportion of PV-positive Ascl1SA6/Bcl2 iNs in layer I was similar to that found in deeper layers (layers II-VI), suggesting that PV-positive iNs are generated similarly across different layers of the cortex (Fig. 5C). Moreover, consistent with the presence of PV-positive iNs, we observed FS properties recorded from Ascl1SA6/Bcl2 iNs located in layer I (Fig. 5D, E). This data suggests that overexpression of Ascl1SA6 together with Bcl2 can induce iNs with hallmarks of FS/PV interneurons independently of layer-specific instructive signals provided by the local environment.

**Fig. 5.**
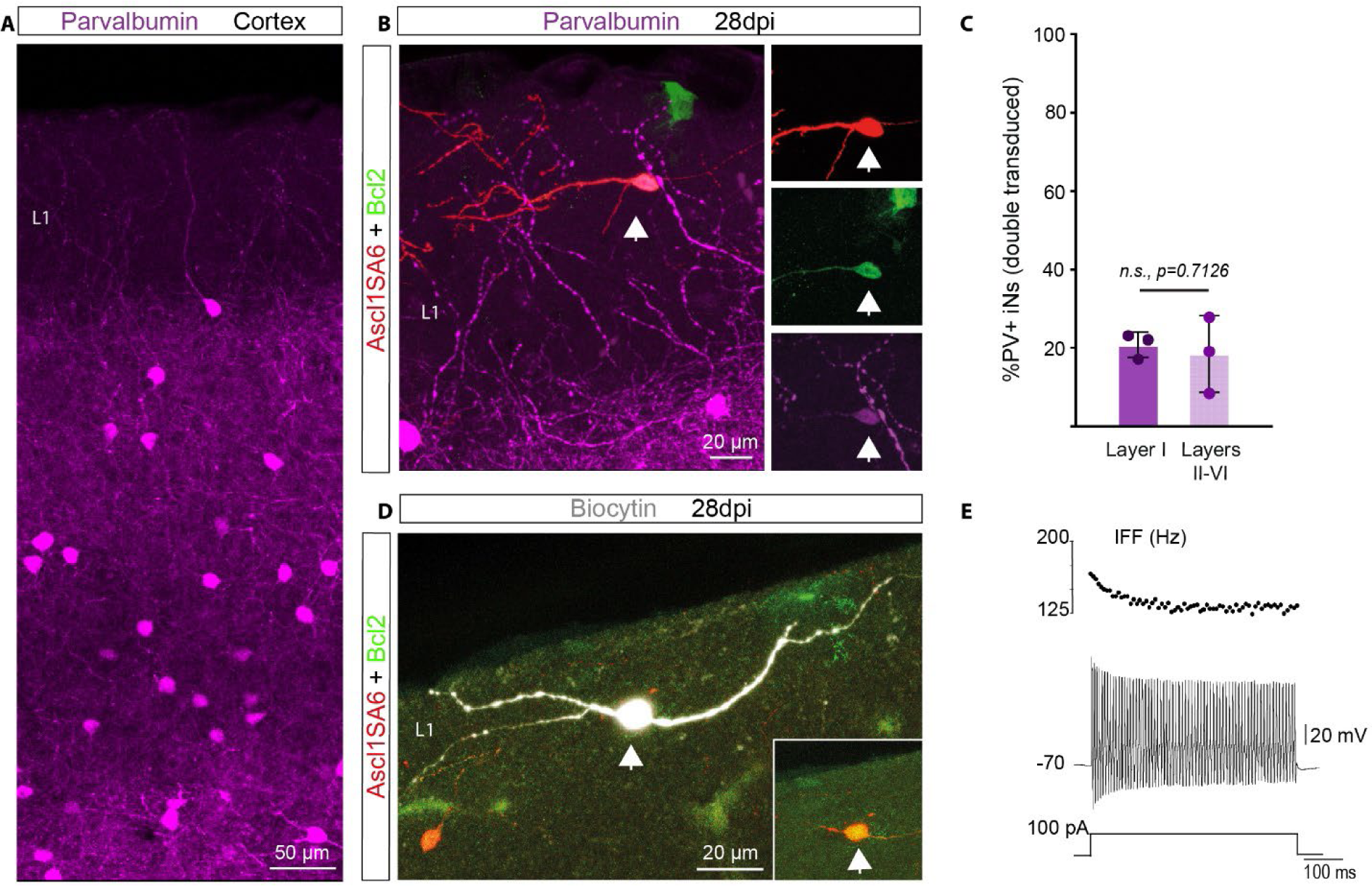
Fast-spiking and parvalbumin-expressing iNs can be located in cortical layer I. (A) Expression of parvalbumin (PV) in the cerebral cortex. Note the absence of PV-positive interneurons in cortical layer I. (B) PV-positive Ascl1SA6/Bcl2 iN (arrow) generated in cortical layer I. (C) Quantification of the percentage of PV positive iNs in layer I and the other layers (II-VI). 20.73 ± 3.2% in layer I, 18.39 ± 9.7% in layers II-VI. 161 PV-iNs, 42 PV+ iNs, n = 3 mice. Data shown as mean ± SD (D) Biocytin-filled Ascl1SA6/Bcl2 iN recorded in layer I. Inset shows the expression of the retroviral reporter proteins DsRed and GFP. (E) Fast-spiking properties (instantaneous firing frequency (IFF) >100 Hz) observed in a Ascl1SA6/Bcl2 iN located in cortical layer I (6 out of 8 FS-iNs). Two-tailed Student’s unpaired t-test in (C).

## Discussion

We show here that neuronal reprogramming of glia in the early postnatal cerebral cortex by the transcription factor Ascl1 is enhanced by using the phospho-site variant Ascl1SA6, in which six serine residues are mutated to alanine (*26*). Stringent genetic fate mapping showed that astrocytes are the main endogenous starting population of the iNs, while very few iNs are derived from OPCs. Moreover, overexpression of Ascl1SA6 in combination with Bcl2 (*6*) not only boosts the efficiency of glia-to-neuron conversion and/or iN survival but also generates iNs with specific histochemical and electrophysiological hallmarks of FS PV-positive interneurons. Finally, we show that FS and PV-expressing iNs can be generated in an ectopic location in the cortex.

We provide compelling evidence that glia-to-neuron conversion is authentic: (1) consistent with our experimental approach that aimed at selectively targeting glia undergoing proliferative expansion we found that iNs effectively incorporated BrdU, when administrated around the time of retrovirus injection; (2) genetic fate mapping experiments demonstrated that approximately 80% of Ascl1SA6/Bcl2 iNs were lineage-traced to astrocytes. While the proportion of astrocytes targeted by the Aldh1l1-CreERT2 mouse line is very high (*34*), it does not capture all cortical astrocytes. It is thus plausible that some of the remaining 20% of non-lineage traced iNs originated from astrocytes that failed to undergo recombination. Overall, these data argue strongly against unspecific labeling of endogenous neurons (*1*, *10*, *15*).

Our results demonstrate the important role of the phosphorylation state of Ascl1 in its neurogenic activity for glia-to-neuron conversion *in vivo*. This is in line with previous findings demonstrating the impact of phosphorylation on proneural factors, as well as studies using similar phosphosite-deficient constructs in progenitors during development and in human neuroblastoma cells (*20*, *26*, *27*, *45*). The enhanced neurogenic potential of the mutant variant reported here could be attributed to different mechanisms of action: i) Higher protein stability of Ascl1SA6 due to differences in ubiquitination (*46*) that might increase the efficiency to target binding sites with less accessibility, such as neuronal differentiation genes; ii) changes in binding properties, e.g., different binding partners by Ascl1SA6 compared to wt Ascl1. Indeed, GSK3-mediated phosphorylation of the proneural factor Neurogenin 2 (Neurog2) promotes binding to LIM-homeodomain transcription factors in the embryonic spinal cord to induce a motor neuron fate (*47*); iii) mutations might cause direct changes in protein structure, enhancing the pioneer activity of Ascl1SA6, especially given that phosphorylation adds a negative charge that might interfere with interactions with negatively charged DNA. Finally, we also confirmed a synergistic effect of Bcl2 to promote neuronal reprogramming and/or enhance the survival of iNs, consistent with previous reports *in vitro* and *in vivo* in combination with the proneural factor Neurog2 (*6*).

Importantly, our results highlight the importance of cellular context for glia-to-neuron conversion. While our retroviral approach targets both proliferative astrocytes and OPCs (*23*), we found that astrocytes are more efficiently converted into iNs by Ascl1SA6/Bcl2 compared to OPCs. Which subtypes of astrocytes (*48*) were converted into iNs and whether astrocyte heterogeneity may have an impact on reprogramming competence remains to be elucidated. Given that OPCs can be transduced by control retroviruses in a proportion of around 20% of the total transduced cells, it is intriguing that less than 3% of Ascl1SA6/Bcl2-iNs were OPC-derived. One intriguing possibility may consist in differential phosphorylation of serine 150 in the second helix region of Ascl1, due to enhanced activity of a non-proline directed serine/threonine kinase in OPCs as compared to astrocytes. Phosphorylation of this serine has been proposed as a general off-switch for proneural factor activity (*49*). Alternatively, neurogenic activity of Ascl1SA6 might induce enhanced cell death of OPCs despite retroviral Bcl2 expression. The limited reprogramming competence of OPCs in the early postnatal cortex contrasts with the high reprogramming competence of OPCs previously observed in the injured adult brain and spinal cord (*5*, *50*, *51*).

A key finding of our study is the observation that Ascl1SA6 promotes the emergence of FS phenotype and PV expression from glia *in vivo,* properties that are hallmarks of FS/PV+ GABAergic interneurons. Dysfunction of these interneurons is implicated in neuropsychiatric disorders (*52*, *53*). Significant efforts have been dedicated to generating this interneuron subtype from induced pluripotent stem cells for disease modeling and potential cell transplantation but has proved challenging (*54–56*). Our data shows that around 20% of Ascl1SA6/Bcl2 iNs express PV protein and 50% among the recorded cells exhibited a FS phenotype. Although our reprogramming approach is unlikely to generate *bona fide* FS/PV+ interneurons, our results suggest that Ascl1SA6/Bcl2 activates gene expression modules contributing to FS/PV identity, including expression of the potassium channel Kv3.1. Interestingly, the activation of such modules can occur in an ectopic location, e.g., cortical layer I, suggesting that transcription factors can exert a determinant effect on the expression of interneuron subclass hallmarks irrespective of influences from the local environment. This does not exclude the possibility that cortical projection neurons could further refine iN properties and distribution, as occurs with endogenous interneurons (*57*).

Taken together, by employing Ascl1SA6 as a reprogramming factor, we succeeded in generating iNs with hallmarks of FS/PV-expressing interneurons from astrocytes *in vivo*. Our findings may thus pave the way towards leveraging lineage reprogramming of glia into subtype specific neurons for restoration of diseased brain circuits.

## Materials and Methods

### Plasmids and retroviruses

Moloney Murine Leukaemia Virus (MMLV)-based retroviral vectors (*58*) were produced to express Ascl1 and Ascl1SA6 as previously described (*23*). Briefly, to generate the pCAG-Ascl1-IRES-DsRed and pCAG-Ascl1SA6-IRES-DsRed retroviral constructs, a cassette containing the coding sequences flanked by attL recombination sites was generated through the excision of the coding sequences for Ascl1 and Ascl1SA6 from pCIG2 parental vectors (*26*). We used the previously described construct for Bcl2: pMIG_Bcl2_IRES_GFP (*6*). We generated a retroviral backbone allowing for polycistronic expression of Ascl1 or Ascl1SA6 and Bcl2 (connected via a T2A peptide sequence) together with DsRed, and under control of the CAG promoter: pCAG-Ascl1-T2A-Bcl2-IRES-DsRed and RV-pCAG-Ascl1SA6-T2A-Bcl2-IRES-DsRed. Viral particles were produced using gpg helper free packaging cells to generate Vesicular Stomatitis Virus Glycoprotein (VSV-G)-pseudotyped retroviral particles (*59*). Retroviral particles were harvested and concentrated from supernatants of transfected packaging cells by ultracentrifugation following standard protocols, re-suspended in TBS (Tris-buffered saline), and stored at −80°C until use. Viral titres used for experiments were in the range of 10^6^-10^8^ transducing units/ml.

### Animals and animal procedures

The study was performed in accordance with the guidelines of the German Animal Welfare Act, the European Directive 2010/63/EU for the protection of animals used for scientific purposes and the Animal (Scientific Procedures) Act 1986 and was approved by local authorities (Rhineland-Palatinate State Authority, permit number 23 177 07-G15-1-031; ethical committee of King’s College London and the UK Home Office, permits numbers PD025E9BC and PP8849003). Mice were kept in a 12:12 h light-dark cycle with food and water *ad libitum*. Male and female C57BL/6J pups were purchased with their mother from Janvier Labs (Le Genest-Saint-Isle, France) or bred in-house from adult mice purchased from Charles River Laboratories (Walden, UK). Male and female transgenic mice used in this study were generated in-house. For this purpose, mice in which the expression of Cre recombinase is driven by mouse GFAP promoter (mGFAP-Cre) (B6.Cg-Tg(Gfap-cre)77.6Mvs/2J, JAX024098) (*35*) or in which tamoxifen-inducible Cre recombinase is driven by the aldehyde dehydrogenase 1 family member L1 locus (Aldh1l1) (Aldh1l1-Cre/ERT2, JAX031008) (*34*) or the mouse NG2 promoter (NG2-CreER^TM^, JAX008538) (*36*) were crossed with an EGFP reporter mouse line (RCE:loxP, JAX032037) (*60*) to generate double transgenic animals (GFAP-Cre/RCE:loxP, Aldh1l1-Cre/ERT2/RCE:loxP or NG2-CreERT2/RCE:loxP). In Aldh1l1-Cre/ERT2 or NG2-iCreERT2/RCE:loxP mice, tamoxifen induction of Cre recombinase activity was performed by daily subcutaneous administration of tamoxifen to pups from P2 to P5 or at P2 and P5, respectively (60 μl each of a 6 mg/ml tamoxifen solution (ApexBio Technology, #B5965) prepared in corn oil (Sigma, Merck, Germany, #C8267) and 10% ethanol). Retroviral injections targeted to the somatosensory and visual cortical areas were performed as previously described (*23*). Briefly, pups at P5 were anesthetized and received a volume of 0.5–1 µl of retroviral suspension using glass capillaries through a small skull incision. After injection, the wound was closed with surgical glue, and pups were left to recover on a warm plate/chamber (37°C) before returning them to their mother. The recovery state was checked daily for a week after the surgery.

### Immunohistochemistry

Tissue preparation and immunostainings were performed as described previously (*23*). Briefly, animals were lethally anesthetized by intraperitoneal administration of 120 mg/kg Ketamine (Zoetis) and 16 mg/kg Xylazine (Bayer) or 1 mg/kg Medetomidine (Orion Pharma) prepared in 0.9% NaCl, and transcardially perfused with 0.9% NaCl followed by 4% paraformaldehyde (PFA, Sigma, Merck, Germany, P6148). The brains were harvested and post-fixed overnight in 4% PFA at 4°C. Then, 40 µm thick coronal sections were prepared using a vibratome (Microm HM650V, Thermo Scientific, Waltham, MA, USA, or Leica VT1000S) and stored at −20°C in a cryoprotective solution containing 20% glucose (Sigma, Merck, Germany, G8270), 40% ethylene glycol (Sigma, Merck, Germany, 324558), and 0.025% sodium azide (Sigma, Merck, Germany, S2202). Using a free-floating procedure, brain sections were washed three times for 15 min with 1X TBS and then incubated for 1.5 h in blocking solution: 2.5% Donkey Serum (Sigma, Merck, Germany, S30); 2.5% Goat Serum (Sigma, Merck, Germany, S26); 0.3% Triton X-100; 1X TBS. Slices were then incubated with primary antibodies diluted in blocking solution for 2–3 h at RT, followed by overnight incubation at 4°C. After three washing steps with 1X TBS, slices were incubated with secondary antibodies diluted in blocking solution for 2 h at RT. Slices were washed twice with 1X TBS, incubated with DAPI 5 µM in 1X TBS for 5 min at RT, and washed three times with 1X TBS. For mounting, slices were washed two times with 1X Phosphate Buffer (PB) and were dried on Superfrost (Thermo Fisher Scientific, Waltham, MA, USA) microscope slides. Sections were further dehydrated with toluene and covered with cover-glasses mounted with DPX mountant for histology (Sigma, Merck, Germany, 06522) or directly mounted with ProlongTMGold (Thermo Fisher Scientific, Waltham, MA, USA, P36930) or with Mowiol (Polysciences, Warrington, PA, USA, #17951-500) supplemented with DABCO (#15154-500, Polysciences, Warrington, PA, USA). The following primary antibodies were used: anti-Doublecortin (Dcx, goat, 1:250, Santa Cruz Biotechnology, Dallas, TX, USA, sc-8066); anti-Doublecortin (Dcx, guinea pig, 1:500, Merck Millipore, Burlington, MA, USA, AB2253); anti-Gamma-aminobutyric acid (GABA, rabbit, 1:300, Sigma, Merck, Germany, A2052); anti-Green Fluorescent Protein (GFP, chicken, 1:1,000, AvesLab, Davis, CA, USA, GFP-1020); anti-Green Fluorescent Protein (GFP, goat, 1:500, Abcam, Cambridge, UK, ab5450); anti-Kv3.1 (mouse IgG1, 1:400, NeuroMab, Davis, CA, USA, 75-041); anti-mCherry (chicken, 1:300, EnCor Biotechnology, Gainsville, FL, USA, CPCA-mCherry); anti-Neuronal Nuclei (NeuN, mouse IgG1, 1:500, Merck Millipore, Burlington, MA, USA, MAB377); anti-Parvalbumin (PV, guinea pig, 1:1000, Synaptic Systems, Göttingen, Germany, 195004); anti-Red Fluorescent Protein (RFP, rabbit, 1:500, Biomol, Hamburg, Germany, 600401379S); anti-Sox9 (mouse IgG1, 1:400, eBiocience, San Diego, CA, USA, GMPR9). Secondary antibodies were made in donkey or goat and conjugated with: A488 (anti-chicken, 1:500, Jackson Immunoresearch, Ely, UK, 703-545-155); A488 (anti-goat, 1:200, Abcam, Cambridge, UK, ab150129); A488 (anti-mouse, 1:200, Thermo Fisher Scientific, Waltham, MA, USA, A21202); A568 (anti-rabbit, 1:500, Thermo Fisher Scientific, Waltham, MA, USA, A11011); A647 (anti-rabbit, 1:500, Thermo Fisher Scientific, Waltham, MA, USA, A31573); A647 (anti-mouse, 1:500, Thermo Fisher Scientific, Waltham, MA, USA, A31571); Cy3 (anti-chicken, 1:500, Dianova, Hamburg, Germany, 703-165-155); Cy3 (anti-rabbit, 1:500, Dianova, Hamburg, Germany, 711-165-152); and Cy5 (anti-goat, 1:500, Dianova, Hamburg, Germany, 705-175-147); Cy5 (anti-guinea pig, 1:500, Jackson Immunoresearch, Ely, UK, 706-175-148).

### Bromodeoxyuridine (BrdU) experiments

BrdU (Sigma, Merck, Germany, #B5002) was administered during the retrovirus delivery and on the following day by intraperitoneal injection at a dose of 50 mg/kg (in 0.9 % NaCl and 0.1 M NaOH). Analysis was performed at 4 weeks post injection. The following modifications were made to the immunohistochemistry protocol for BrdU staining: sections were pretreated with 2N HCl and 0.5% Triton X-100 for 30 min at 37°C followed by neutralization with sodium-tetraborate (Na2B4O7, 0.1M, pH 8.6) for 30 min at RT before incubation with blocking solution consisting of 3% bovine serum albumin (Sigma, Merck, Germany, #A9418) and 0.3% Triton X-100 in PBS 1X. This blocking solution was also used for dilution of the primary and secondary antibodies: rat anti-BrdU (1:200; Serotec, OBT0030) and donkey anti-rat 647 (1:500, Jackson Immunoresearch, Ely, UK, 712-606-150).

### Single-molecule fluorescence *in situ* hybridization (smFISH)

Single-molecule fluorescent in situ hybridization with RNAscope technology was performed using the Multiplex Fluorescent V2 kit (Advanced Cell Diagnostics, ACD) according to the manufacturer instructions. All solutions were prepared in RNase-free Diethyl pyrocarbonate (DEPC)-treated water (Sigma, Merck, Germany, D5758). Hybridizations towards targets were performed with probes Pvalb-C1 (#421931), Sst-C3 (#404631) and Kcnc1-C1 (#564521), and control hybridizations were performed with a positive control probe (PN320881) and a negative control probe (PN320871). Sections were placed in 0.05M TBS and mounted in Phosphate Buffer (PB) 0.1M on Superfrost slides (J1800AMNZ, Menzel). Sections were let to dry overnight at RT and then at 40°C for 40 min in HybEZII oven (ACD). They were then rinsed with distilled water (dH_2_0) and dehydrated with 50%, 70% and 100% ethanol (EtOH), 5 minutes each. Then, sections were treated with hydrogen peroxide (Multiplex Fluorescent V2 kit, ACD) for 10 min at room temperature (RT), washed 3 x 3-5 min with dH_2_O followed by 3 x 3-5 min in TBS 1x before incubation with antigen retrieval solution (Multiplex Fluorescent V2 kit, ACD) for 10-15 min at 90°C. Sections were rinsed with dH_2_O, dipped in EtOH 100%, and air dried for approximately 5 min. They were then incubated with Protease III (Multiplex Fluorescent V2 kit, ACD) for 10-15 min at 40°C in HybEZII oven. The sections were washed 3 x 3-5 min in dH_2_O, incubated for 2h with the probes at 40°C in HybEZII oven, then washed 3 x 3-5 min in wash buffer (Multiplex Fluorescent V2 kit, ACD), and kept overnight in 5X Saline Sodium Citrate (SSC, prepared according to ACD’s user manual). On the next day, the sections were successively incubated with AMP1 (Multiplex Fluorescent V2 kit, ACD) for 30 min at 40°C in HybEZII oven, AMP2 (Multiplex Fluorescent V2 kit, ACD) for 30 min at 40°C in HybEZII oven and AMP3 (Multiplex Fluorescent V2 kit, ACD) for 15 min at 40°C in HybEZII oven, with intermediate washes of 3 x 3-5 min in Wash Buffer (ACD). The signal was then developed by incubation with HRP-C1 or HRP-C3 (adapted to the channel of the probe, Multiplex Fluorescent V2 kit, ACD) for 15 min at 40°C in HybEZII oven, followed by Opal dye 690 diluted in TSA buffer (1:1000, #322809, ACD) for 30 min at 40°C in HybEZII oven and then HRP-blocker (Multiplex Fluorescent V2 kit, ACD) for 15 min at 40°C in HybEZII oven, with intermediate washes of 3 x 3-5 min in Wash Buffer (ACD). Then, the sections were incubated with DAPI 5 µM in 1X TBS for 7 min at RT and transferred into TBS 0.1M for further processed for immunostaining. After permeabilization with PBS 0,25% Triton for 20 min at RT, sections were incubated with blocking solution consisting of 0,3% Triton, 3% BSA (Bovine Serum Albumin, Scientific Laboratory Supplies, #A2153), 10% donkey or goat serum (or 5% donkey and 5% goat serum) in PBS 1X for 2h at RT, and then incubated with primary antibodies prepared in 0,3% Triton, 3% BSA, 5% Serum in PBS 1X overnight+24h at 4°C. They were washed 3 times in PBS 1X and incubated with secondary antibodies for 2h. Finally, they were transferred to 0.1M phosphate buffer (PB) and mounted with ProlongTMGold (Thermo Fisher Scientific, Waltham, MA, USA, P36930). The following primary antibodies were used: chicken anti-GFP: 1:200; rabbit anti-RFP: 1:100. Secondary antibodies were made in donkey or goat and conjugated with: A488 (anti-chicken, 1:200), A568 (anti-rabbit, 1:250).

### Confocal microscopy and quantifications

Immunostainings and RNAscope signals were imaged with a TCS SP5 (Leica Microsystems, Wetzlar, Germany) confocal microscope (Institute of Molecular Biology, Mainz, Germany) equipped with four PMTs, four lasers (405 Diode, Argon, HeNe 543, HeNe 633) and a fast-resonant scanner using a 20X dry objective (NA 0.7) or a 40X oil objective (NA 1.3), or with a Zeiss LSM 800 confocal microscope (Carl Zeiss Microscopy, Jena, Germany) equipped with four solid-state lasers (405, 488, 561, and 633 nm) at a 20X (NA 0.8) or 40X (NA 1.3) objectives (Centre for Developmental Neurobiology, King’s College London), or with a Zeiss Axio Imager.M2 fluorescent microscope equipped with an ApoTome (Carl Zeiss Microscopy, Jena, Germany) at a 20x dry objective (NA 0.7) or a 63x oil objective (NA 1.25). Serial Z-stacks spaced at 0.3–2.13 mm distance were acquired to image the whole thickness of the brain sections. Images were captured with Leica Application Suite or Zeiss ZEN softwares. For the figures, maximum intensity projections from the Z-stacks were generated using the function provided by the software. Cell quantifications were performed using ZEN software or ImageJ 1.51v software (National Institute of Health, USA). Cell counts were done by navigating through the Z-stacks of images, allowing the accurate visualization of each transduced cell. For fate-mapping experiments, we counted the number of 1) RFP/EGFP/Dcx or NeuN-triple-positive cells (i.e., fate-mapped induced neurons), 2) RFP/Dcx or RFP/NeuN-double positive induced neurons, 3) RFP/EGFP-double positive transduced cells and 4) RFP only positive transduced cells. Cells in each group are expressed as a percentage of the total number of reporter- or double-reporter-positive transduced cells. Results are expressed as mean ± SD. Cells were quantified from at least 3-5 sections from three independent mice (minimum nine sections in total).

For RNAscope signal, the DsRed reporter signal from transduced cells or the DAPI+ nuclei from endogenous neurons were used to draw an ROI. The number of mRNA transcripts was manually counted by navigating through the Z-stacks of confocal images obtained with a 40X (NA 1.3) objective and spaced at 0.5-0.75 µm distance. The number of mRNA transcripts was expressed as total number of dots per DsRed+ cell or endogenous neuron. For each experiment, quantification was done in cells from three independent mice.

### Electrophysiological recordings

#### Slice preparation

Mice anesthetized with isoflurane (Forane, Abbvie, Illinois, USA) were decapitated and brains were removed into chilled artificial cerebro-spinal fluid (ACSF) solution containing the following (in mM): NaCl, 85; Sucrose, 73; KCl, 2.5; NaHCO_3_, 25; CaCl_2_, 0.5; MgCl_2_, 7; NaH_2_PO_4_, 1.25 and glucose 10; saturated with 5% CO_2_ and 95% O_2_, pH 7.4. Coronal 300 μm-thick cortical slices were cut using a vibratome (VT1200 S, Leica, Wetzlar, Germany) and transferred to standard ACSF (34°C, 10-15 min) containing (mM): NaCl, 125; KCl, 2.5; NaHCO_3_, 25; CaCl_2_, 2; MgCl_2_, 1; NaH_2_PO_4_, 1.25 and glucose 12; pH 7.4. Subsequently, the slices were stored in standard ACSF at room temperature (21 ± 2 °C) for at least 1 h before experiments started. Individual slices were placed in a recording chamber, superfused (1-2 ml min−1, standard ACSF) and mounted on an upright microscope (Axio Imager 2, Zeiss, Jena, Germany or Slice Scope Pro 6000 System, Scientifica, Uckfield, UK). Cells were visualized with oblique contrast or Dodt gradient contrast optics using a 40X (0.70 numerical aperture) water immersion objective and with a Hamamatsu Orca-Flash 4.0 camera (Hamamatsu, Japan). For the identification of the retrovirally transduced cells, GFP and DsRed fluorescence was revealed by LED (CoolLED pE-100, Andover, UK) excitation with appropriate excitation and emission filters.

#### Electrophysiology

Patch-clamp whole-cell recordings were performed at 30 ± 2 °C with an in-line heater (Scientifica, Uckfield, UK). Recording pipettes (10-15 MΩ) were pulled from borosilicate capillary glass (BF150-86-10, Sutter Instruments, Novato, USA or 1B150F-4, World Precision Instruments, Sarasota, USA) in a horizontal pipette puller (P-1000 Micropipette puller, Sutter Instruments, Novato, USA) and were filled with (in mM): K-gluconate, 125; NaCl, 5; Na_2_-ATP, 2; MgCl_2_, 2; EGTA, 1; HEPES, 10; and biocytin, 10 to allow a posterior morphological analysis; pH 7.4, osmolarity, 280 mOsm. Voltage and current clamp recordings were obtained using Axopatch 200B or MultiClamp 700B amplifier (Molecular Devices, San Jose, USA), digitized (Digidata 1440A or 1550B, Molecular Devices, San Jose, USA), and acquired at 20 kHz onto a personal computer using the pClamp 10 software (Molecular Devices, San Jose, USA) which was also used for further analysis. Criteria to include cells in the analysis were: 1) visual confirmation of GFP or DsRED in the pipette tip, 2) attachment of the labeled soma to the pipette when suction was performed, 3) seal resistances between 4 and 18 GΩ, 4) initial series resistance less than 40 MΩ and did not change by more than 20% throughout the recording period. Whole-cell capacitance and series resistances were not compensated. In current-clamp recordings, the membrane potential was kept at −70 mV or adjusted to different values, if necessary, by passing a holding current. Passive and active membrane properties were recorded by applying a series of hyperpolarizing and depolarizing current steps (2 or 10 pA steps, 500 ms). The instantaneous firing frequency (IFF) was calculated as the inverse value of the inter-spike interval, which was determined by the time difference between adjacent action potential peaks. Maximum firing frequency was calculated dividing the number of spikes by the step duration. The resting membrane potential (RMP) was assessed immediately after break-in by reading the voltage value in the absence of current injection (I=0 configuration). Input resistance (R_in_) was calculated from the peak of the voltage response to a −20 pA, 500 ms current step according to Ohm’s law (Vhold = −70 mV). The membrane capacitance (C_m_) was obtained with the following equation: membrane time constant (τ_m_) = R_in_.C_m_. τ_m_ was derived from single exponential fitted to voltage response to - 20 pA, 500 ms current step. The action potential properties were analyzed on the first spike observed at the rheobase. Spike amplitude was measured from threshold to positive peak and after-hyperpolarization (AHP) amplitude, from threshold to negative peak during repolarization. Spike width was measured at the half amplitude of the spike. Frequency and amplitude of the spontaneous excitatory postsynaptic currents were recorded during 60 s in mode Gap-free at Vhold = −70 mV and the analysis was done using Clampfit template search (Molecular Devices, San Jose, USA). No correction was made for the junction potential between the pipette and the ACSF.

In some experiments, tetrodotoxin (TTX, 1 µM, Tocris, Bio-Techne, Minneapolis, USA) to block Na channels or 6-Cyano-7-nitroquinoxaline-2,3-dione (CNQX, 10 µM, Tocris, Minneapolis, USA) to block AMPA/kainate receptors were introduced to the standard ACSF.

### Morphological identification of the recorded cells

Retroviral transduced cells expressing DsRed or GFP were first imaged in acute slices. During whole-cell patch-clamp recordings, cells were filled with biocytin included in the pipette. Then, slices were fixed by immersion in 4% PFA for 24 hours. Following TBS rinsing, the slices were blocked with 0.5% BSA in TBS (1 hour) and then incubated in TBS containing 0.3% Triton X-100 with the streptavidin-fluorophore complex, 1:400, for 2 hours, and subsequently mounted for confocal imaging.

### Statistical analysis

Statistical analysis was performed using OriginLab (Northampton, Massachusetts, USA) or GraphPad Prism 5 (GraphPad, San Diego, CA, USA) or SPSS Statistics V5 (IBM). Data are represented as means ± SD or with box and whisker plots, which give the median, 25^th^ and 75^th^ percentiles and range. The normality of distribution was assessed using Shapiro-Wilk test. The significance of the differences between groups was analysed by independent *t-test* (for samples with normal distribution) or using the Mann–Whitney U test (for samples without normal distribution). The statistical test used is described in the figure legend. The number of independent experiments (n), number of cells analysed, and the number of cells recorded for electrophysiology are reported in the figure legends.

## Acknowledgments

We are grateful to the members of the Berninger laboratory for their helpful comments and critical feedback over the course of this study. We thank S. Leaman for critical reading of the manuscript. We are grateful to N. Carvajal Garcia for excellent technical support and genotyping. We are grateful to B. Rico (King’s College London) for her support throughout the project. We acknowledge support from the Microscopy Core Facility of the Institute of Molecular Biology (IMB) Mainz.

## Author contributions

Conceptualization: B.B.

Design of experiments: N.M., S.P., A.B.A., B.B.

Performed and analyzed *in vivo* experiments: N.M., S.P., A.B.A., C.G., F.F.S., R.W.

Performed and analyzed RNAscope experiments: S.P., A.B.A.

Electrophysiology experiments: N.M.

Formal analysis: N.M., S.P., A.B.A.

Visualization: N.M., S.P., B.B.

Resources: C.S., M.K., S.G. Supervision: B.B.

Writing—original draft: N.M., S.P., A.B.A, B.B.

Writing—review & editing: all authors

## Data and materials availability

Any reagents that this study generated will be shared by the corresponding author upon reasonable request. All other study data are included in the article and/or Supplementary material.

---

**Fig. S1.**
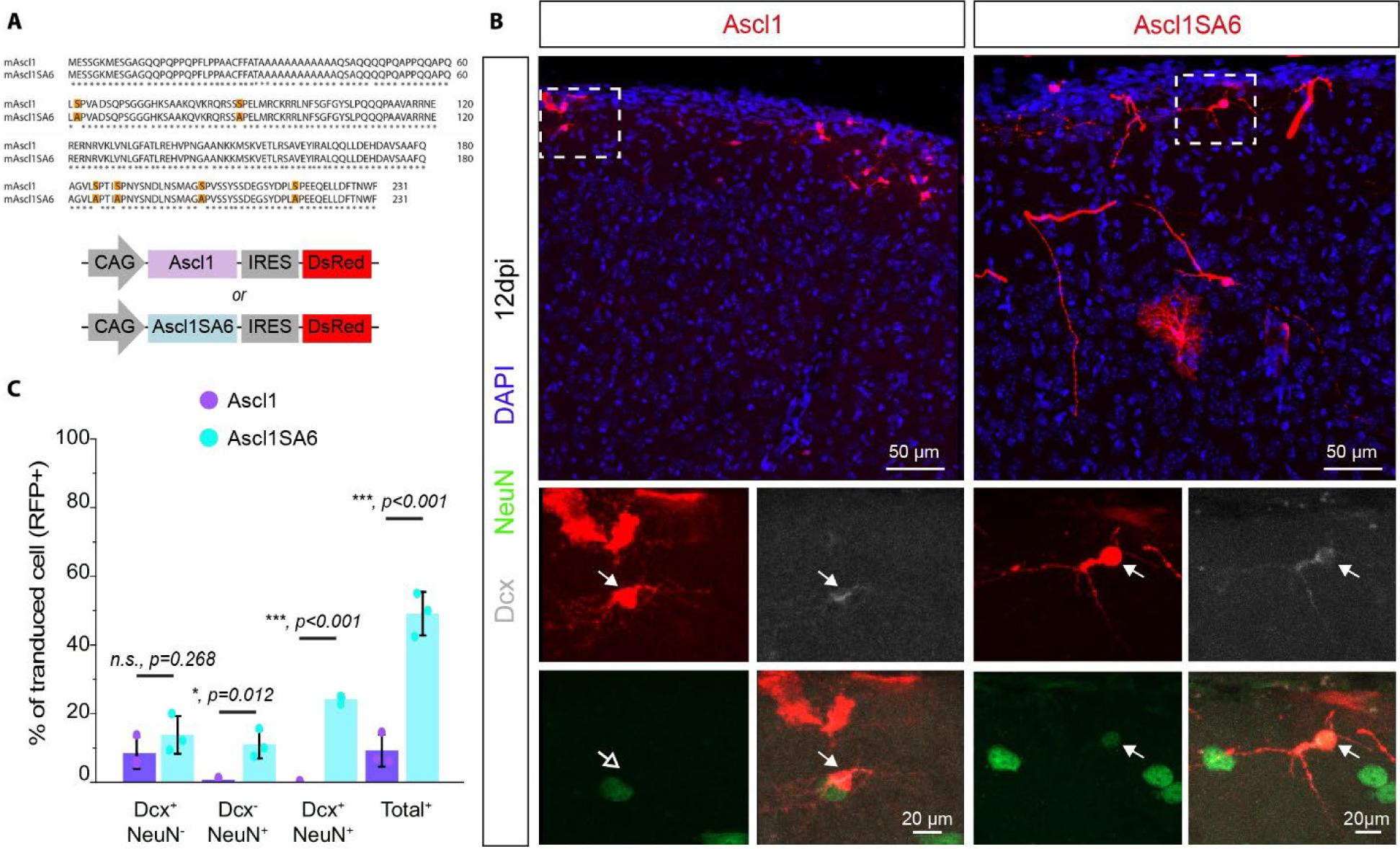
Ascl1SA6 converts postnatal cortical glia into iNs more efficiently than Ascl1 (Related to Fig. 1). (A) Scheme showing the six serine residues mutated to alanine in mouse Ascl1 (mAscl1) to generate the mutant Ascl1SA6 and final expression cassettes in the retroviral constructs. (B) Ascl1- or Ascl1SA6-transduced cells at 12 dpi. High-magnification images from insets (white boxes) depicting presence or absence of Dcx and NeuN expression in transduced cells. Note the maintenance of glial morphology in Ascl1-transduced cells and the acquisition of neuronal morphology in Ascl1SA6 iNs. Empty arrows indicate marker-negative cells. (C) Proportion of transduced cells expressing Dcx, NeuN or both neuronal markers at 12 dpi. 8.5 ± 4.6% Dcx only, 0.7 ± 0.6% NeuN only, 0.0 ± 0.0% Dcx and NeuN, 341 cells, n = 3 mice for Ascl1; 13.8 ± 5.5% Dcx only, 11.1 ± 4.1% NeuN only, 24.2 ± 1.3% Dcx and NeuN, 118 cells, n = 3 animals for Ascl1SA6. Data shown as mean ± SD. Two-tailed Student’s unpaired t-test in (C).

**Fig. S2.**
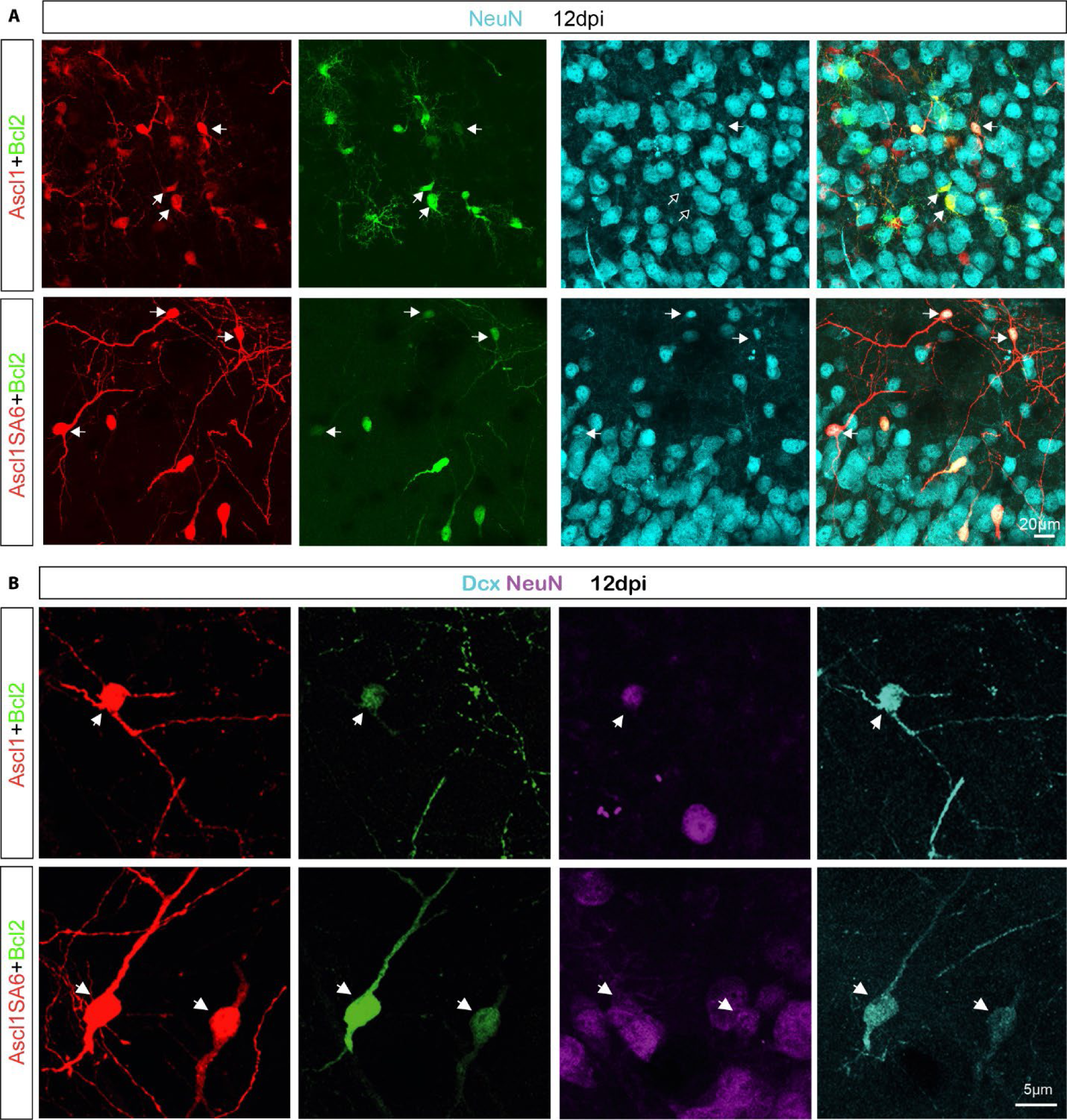
Highly efficient conversion of glia into neurons by Ascl1SA6 and Bcl2 (Related to Fig.1). (A) High-magnification images depicting NeuN expression in co-transduced cells. Empty arrows indicate marker-negative cells at 12 dpi. (B) High-magnification images depicting Dcx and NeuN expression in co-transduced cells at 12dpi.

**Fig. S3.**
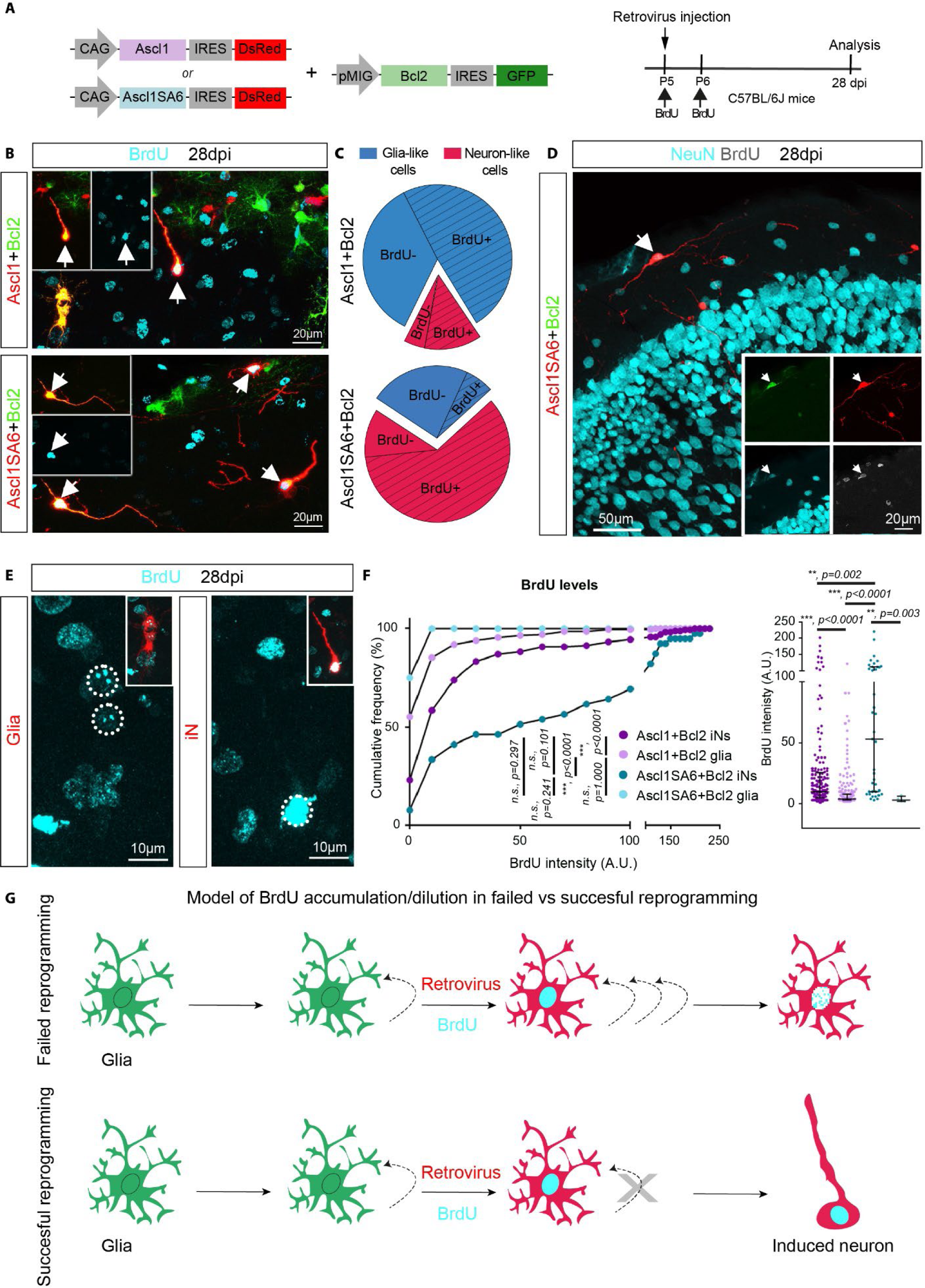
iNs incorporate BrdU administered at the time of retrovirus injection (Related to Fig. 1). (A) Experimental design. (B) BrdU labeling of co-transduced cells that had acquired neuron-like morphology (arrows) at 28 dpi. For better visualization of BrdU labeling, inset shows channel separation for DsRed/GFP and BrdU signals (C) Pie charts showing the proportion of glia-like or neuron-like co-transduced cells that were BrdU-positive (+) at 28 dpi. Notice that the vast majority of co-transduced cells with neuronal morphology were BrdU+ (red-slashed section). 1217 cells, 17 ± 5.3% neuron-like cells, from which 72.9 ± 1.7 % were BrdU+, n = 3 mice for Ascl1/Bcl2; 65 cells, 71 ± 12.5% neuron-like cells, from which 85.8 ± 5.6 % were BrdU+, n = 3 mice for Ascl1SA6/Bcl2. Data shown as mean ± SD. (D) BrdU labeling of NeuN+ iN (arrow). Inset shows channel separation for the signals (E) BrdU-signal levels in transduced glia and iNs. Inset shows channel separation for DsRed/GFP and BrdU signals (F) Graph showing the cumulative frequency of BrdU-signal intensity (left graph) and individual BrdU-signal intensity values (right graph) in transduced cells. Data shown as median with interquartile range. 400 cells analyzed (161 Ascl1/Bcl2-BrdU+ iNs, 196 Ascl1/Bcl2-BrdU+ glia, 39 Ascl1SA6/Bcl2-BrdU+ iNs, 4 Ascl1SA6/Bcl2-BrdU+ glia). (G) Scheme depicting a model of BrdU accumulation/dilution in failed vs successful reprogramming. Kruskal-Wallis for comparison of cumulative intensity frequency distribution in (F, left) and median test with Bonferroni correction for comparison of median intensity values in (F, right).

**Fig. S4.**
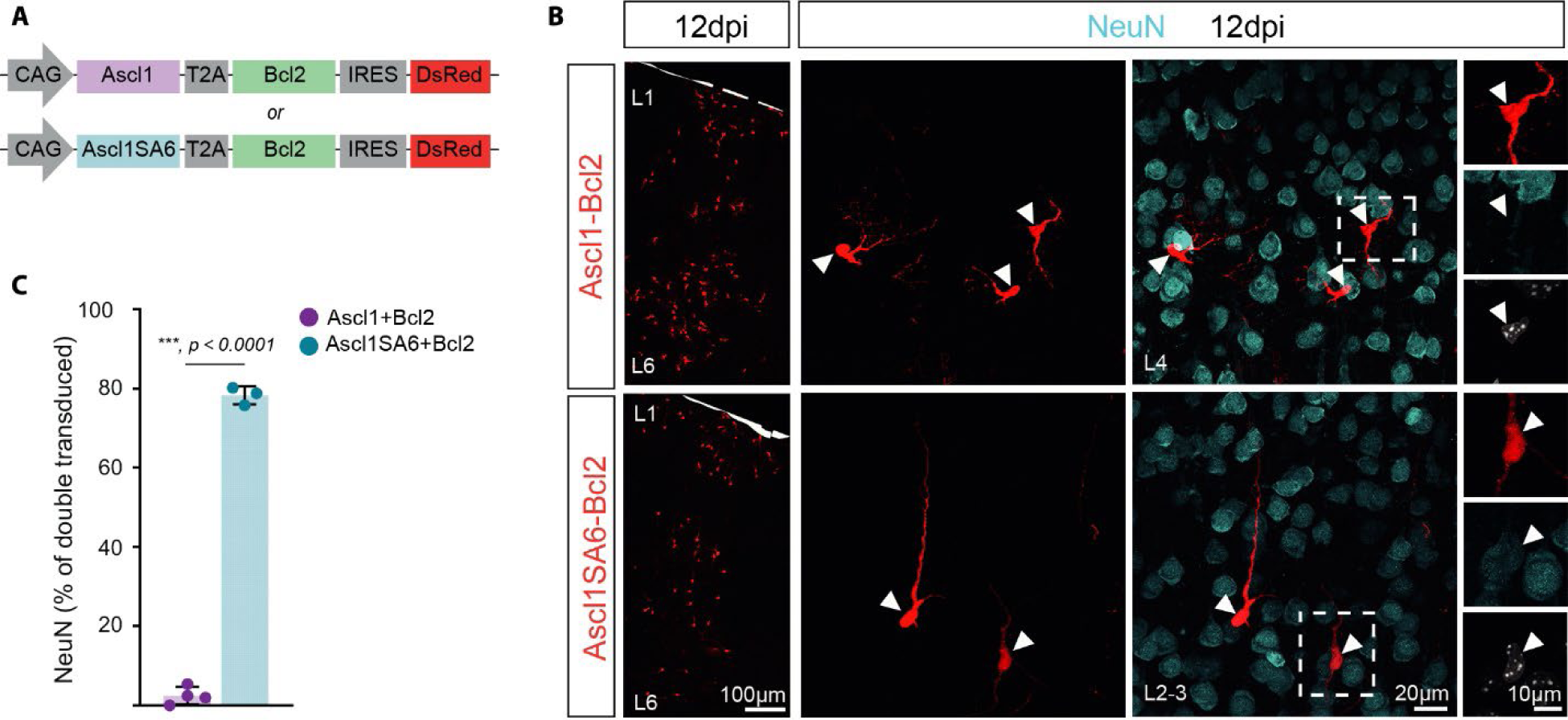
A single retroviral vector encoding for Ascl1SA6-Bcl2 also induces high efficiency of neuronal reprogramming (Related to Fig. 2) (A) Scheme showing the retroviral constructs encoding for both Ascl1 and Bcl2 (Ascl1-Bcl2) or Ascl1SA6 and Bcl2 (Ascl1SA6-Bcl2) in a single construct. (B) NeuN expression in Ascl1-Bcl2 or Ascl1SA6-Bcl2 transduced cells at 12 dpi. The left panels show low-magnification images of the transduced cells. Central panels show high magnification of transduced cells. Cells from insets (white boxes) are shown on the right panels. (C) Quantification of the percentage of transduced cells expressing NeuN at 12 dpi. 7.0 ± 4.4%, 1002 cells, n = 4 mice for Ascl1-Bcl2; 78.3 ± 2.3%, 1224 cells, n = 3 mice for Ascl1SA6-Bcl2. Data shown as mean ± SD. Two-tailed Student’s unpaired t-test in (C).

**Fig. S5.**
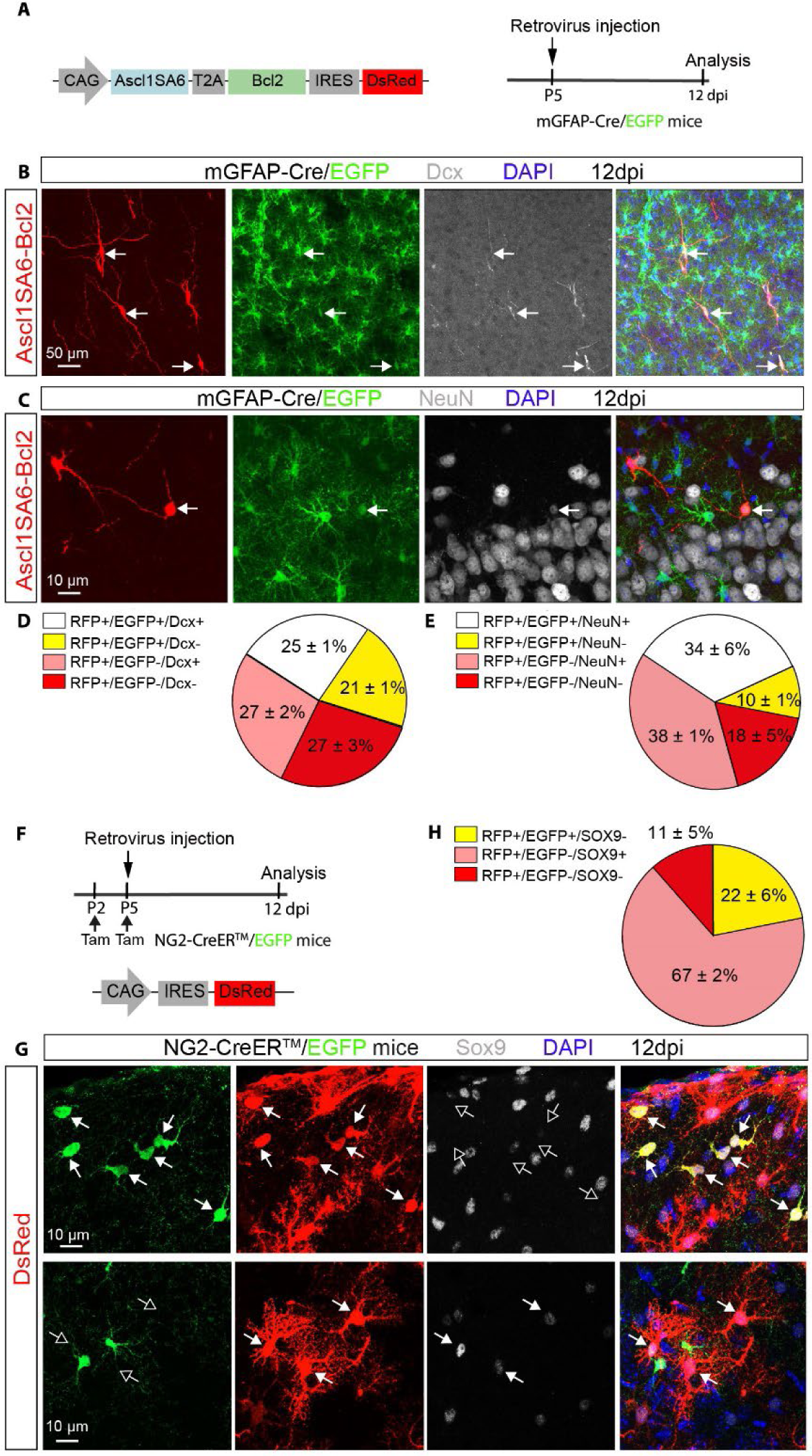
Astroglial origin of the vast majority of Ascl1SA6 and Bcl2 iNs (Related to Fig. 2). (A) Experimental design. (B-C) Ascl1SA6-Bcl2-derived iNs expressing Dcx (B) or NeuN (C) and EGFP (arrows) in the mGFAP-Cre/RCE:loxP mouse line at 12 dpi, demonstrating their astroglial origin. (D-E) Pie charts showing the relative number of transduced cells (DsRed+) co-expressing EGFP and/or Dcx (D), and/or NeuN (E), or DsRed only in mGFAP-Cre/RCE:loxP mice at 12dpi. 503 cells, n = 3 mice for Dcx analysis; 632 cells, n = 3 mice for NeuN analysis. Data shown as mean ± SD. (F) Experimental design. (G) Control-transduced cells co-expressing EGFP and lacking Sox9 expression (upper raw) or vice versa (lower raw) at 12 dpi. Empty arrows indicate marker-negative cells. (H) Pie chart showing the relative number of transduced cells (DsRed+) expressing Sox9, EGFP or DsRed only in NG2CreERTM/RCE:loxP mice at 12dpi. 1374 cells, n = 3 mice. Data shown as mean ± SD.

**Fig. S6.**
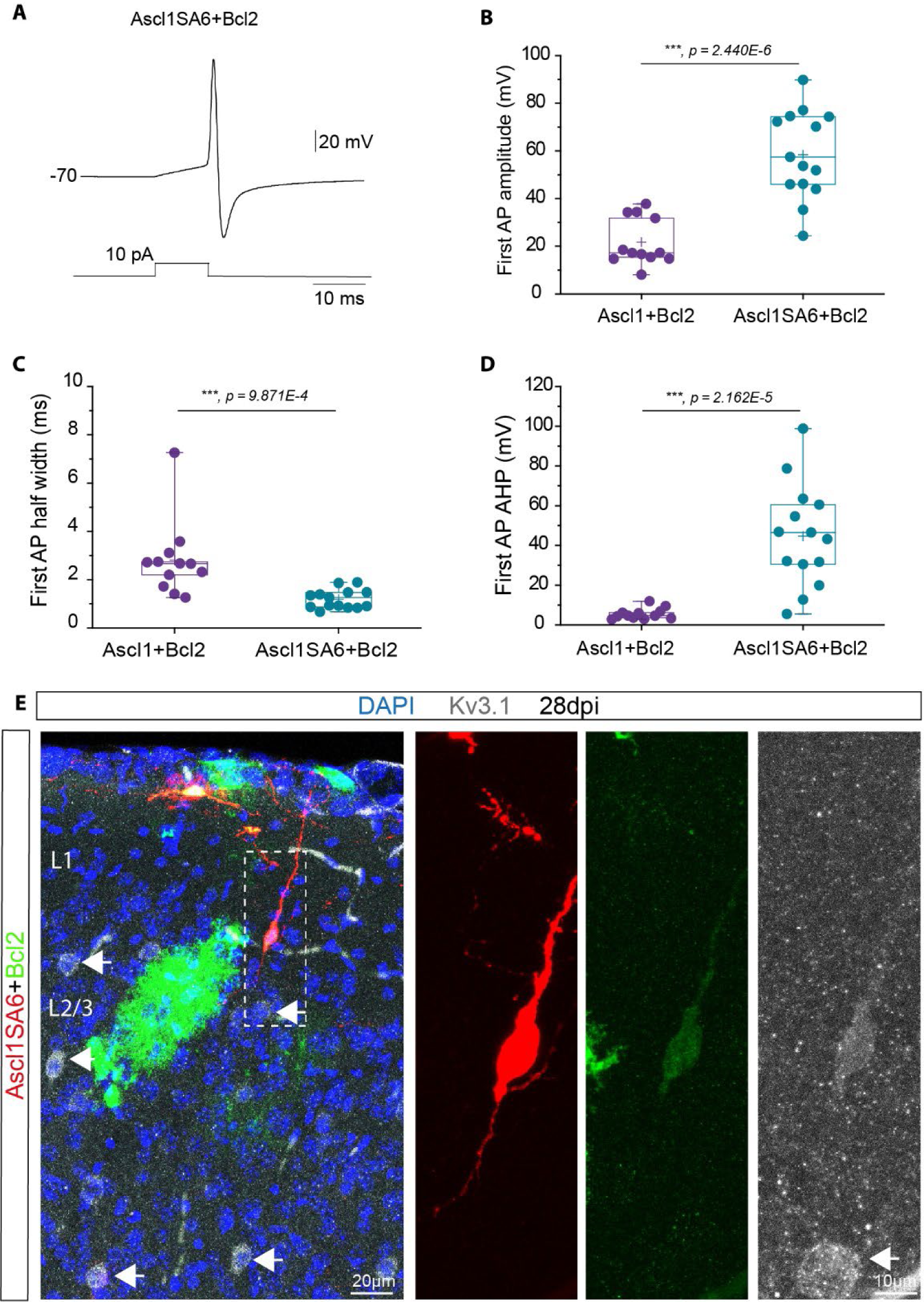
Ascl1SA6/Bcl2 iNs exhibit narrow action potentials with well-developed afterhyperpolarization, and express Kv3.1 (Related to Fig. 4). (A) Action potential recorded from an Ascl1SA6/Bcl2 iN after a brief depolarizing current pulse (10 ms), 28 dpi. (B-D) Action potential (AP) amplitude, AP half-width and AHP amplitude were measured for the first spike generated by Ascl1/Bcl2 (12 cells, n = 5 mice) and Ascl1SA6/Bcl2 (14 cells, n = 8 mice) iNs in response to 500-ms current injection at 28 dpi. Each dot represents one cell. Data shown with box and whisker plots, which give the mean (+), median, 25th and 75th percentiles and range. (E) Kv3.1 protein expression in an Ascl1SA6/Bcl2 iN. Note the high level of Kv3.1 in endogenous interneurons (arrows). High-magnification images correspond to the inset (white box). Two-tailed Student’s unpaired t-test in (B-D).

**Fig. S7.**
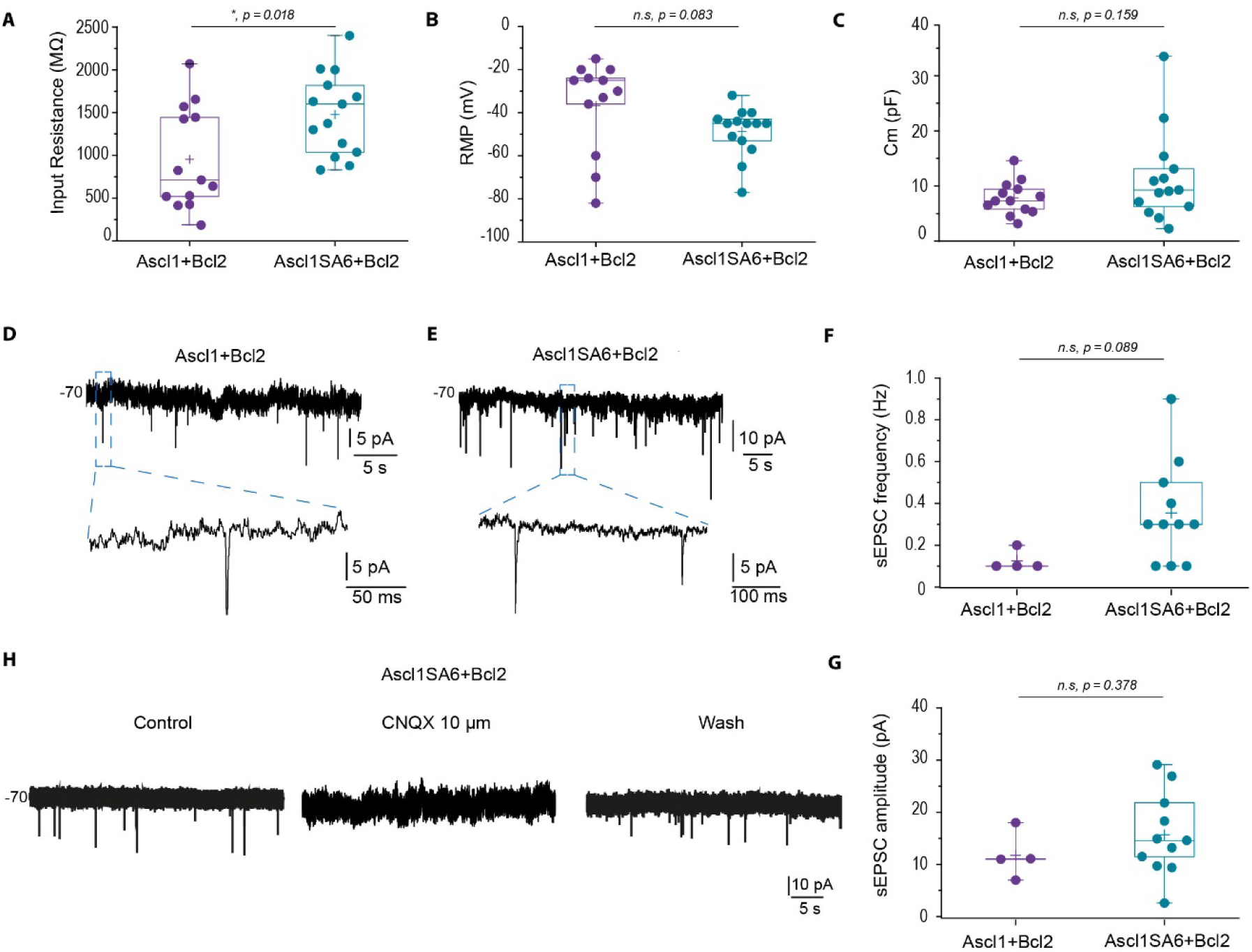
The vast majority of Ascl1SA6/Bcl2 iNs receive excitatory inputs (Related to Fig. 4). (A-C) Input resistance, resting membrane potential (RMP), and membrane capacitance (Cm) of Ascl1/Bcl2 (13 cells, n = 5 mice) and Ascl1SA6/Bcl2 (14 cells, n = 8 mice) iNs at 28 dpi. Each dot represents one cell. Data shown with box and whisker plots, which give the mean (+), median, 25th and 75th percentiles and range. (D-E) Spontaneous synaptic inputs recorded from an Ascl1/Bcl2 iN (D, 4 of 13 cells, n = 5 mice) or an Ascl1SA6/Bcl2 iN (E, 11 of 14 cells, n = 8 mice). Expanded presentations show individual events. (F-G) Frequency and amplitude of spontaneous excitatory postsynaptic currents (sEPSC) recorded from Ascl1/Bcl2 or Ascl1SA6/Bcl2 iNs. 4 out of 13 Ascl1/Bcl2 cells and 11 out of 14 Ascl1SA6/Bcl2 cells exhibited sEPSC and were therefore included in the analysis. Each dot represents one cell. Data shown with box and whisker plots, which give the mean (+), median, 25th and 75th percentiles and range. (H) Synaptic currents in Ascl1SA6/Bcl2 iNs were blocked by the AMPA/kainate receptor blocker CNQX and were recovered after the washout of the drug, confirming their excitatory nature (2 of 2 cells). Two-tailed Student’s unpaired t-test in (A-C, F,G).

